# CCR4 and CCR7 differentially regulate thymocyte localization with distinct outcomes for central tolerance

**DOI:** 10.1101/2022.04.03.486911

**Authors:** Yu Li, Pablo Guaman Tipan, Hilary J. Selden, Jayashree Srinivasan, Laura P. Hale, Lauren I.R. Ehrlich

## Abstract

Central tolerance ensures autoreactive T cells are eliminated or diverted to the regulatory T cell lineage, thus preventing autoimmunity. To undergo central tolerance, thymocytes must enter the medulla to test their TCRs for autoreactivity against the diverse self-antigens displayed by antigen presenting cells (APCs). While CCR7 is known to promote thymocyte medullary entry and negative selection, our previous studies implicate CCR4 in these processes, raising the question of whether CCR4 and CCR7 play distinct or redundant roles in central tolerance. Here, synchronized positive selection assays, 2-photon timelapse microscopy, and quantification of TCR-signaled apoptotic thymocytes, demonstrate that CCR4 and CCR7 promote medullary accumulation and central tolerance of distinct post-positive selection thymocyte subsets. CCR4 is upregulated within hours of positive selection signaling and promotes medullary entry and clonal deletion of immature post-positive selection thymocytes. In contrast, CCR7 is expressed several days later and is required for medullary localization and negative selection of mature thymocytes. In addition, CCR4 and CCR7 differentially enforce self-tolerance, with CCR4 enforcing tolerance to self-antigens presented by activated APCs, which express CCR4 ligands. Our findings show that CCR7 expression is not synonymous with medullary localization and support a revised model of central tolerance in which CCR4 and CCR7 promote early and late stages of negative selection, respectively, via interactions with distinct APC subsets.

## Introduction

Self-tolerance of the T-cell repertoire is established in thymus through the process of central tolerance, which encompasses both negative selection and regulatory T cell (Treg) induction (Klein et al., 2014). To avoid autoimmunity, developing T cells must be broadly tolerized not only to ubiquitously expressed self-antigens, but also to proteins expressed by distinct differentiated cell types. The thymic medulla is a specialized environment in which central tolerance to such diverse self-antigens is enforced. After positive selection in the cortex, thymocytes migrate into the medulla, where they interact with medullary antigen presenting cells (APCs), including thymic dendritic cells (DCs) and medullary thymic epithelial cells (mTECs). Collectively, these APCs enforce self-tolerance by presenting self-peptides from the majority of the proteome on MHC complexes, such that thymocytes expressing autoreactive T cell receptors undergo clonal deletion or diversion to the Treg lineage (Ehrlich, 2016; Klein et al., 2014; Klein et al., 2019; Lancaster et al., 2019). AIRE^+^ mTECs express >80% of the proteome, including *Aire*- dependent tissue restricted antigens (TRAs) that are otherwise expressed in only a few peripheral tissues (Abramson & Anderson, 2017; Anderson & Su, 2016; Mathis & Benoist, 2007; Meredith et al., 2015). DCs also present numerous self-antigens, including those acquired from mTECs, from circulation, or from peripheral tissues and trafficked into the thymus (Atibalentja et al., 2009; Bonasio et al., 2006; Koble & Kyewski, 2009). The importance of inducing thymocyte tolerance to medullary self-antigens is evidenced by multi-organ autoimmunity that ensues in *Aire*-deficient mice and APECED patients (Anderson et al., 2002; Finnish-German, 1997; Nagamine et al., 1997), in whom thymic expression of TRAs is greatly diminished. Failure to express even a single medullary TRA can result in impaired tolerance and subsequent T-cell mediated autoimmune pathology (DeVoss et al., 2006). Notably, individual TRAs are expressed by only 1-3% of *Aire*^+^ mTECs (Brennecke et al., 2015; Meredith et al., 2015; Sansom et al., 2014), resulting in a sparse mosaic of self-antigen display throughout the medulla. Because thymocytes reside in the medulla for only 4-5 days (McCaughtry et al., 2007), tight spatiotemporal regulation is required to ensure that post-positive selection thymocytes encounter the full spectrum of medullary self-antigens required to enforce self-tolerance.

To facilitate medullary entry after positive selection, thymocytes upregulate chemokine receptors that promote their directional migration towards medullary-biased chemokine gradients (Bleul & Boehm, 2000; Hu, Lancaster, & Ehrlich, 2015; Lancaster et al., 2018; Petrie & Zuniga-Pflucker, 2007). Notably, CCR7 has been shown to play a critical role in directing chemotaxis of post-positive selection thymocytes towards the medulla, where CCR7 ligands are expressed, enhancing thymocyte accumulation within the medulla, enforcing negative selection to TRAs, and averting autoimmunity (Ehrlich et al., 2009; Kozai et al., 2017; Kurobe et al., 2006; Nitta et al., 2009; Ueno et al., 2004). Given the significance of CCR7 in promoting thymocyte medullary entry, CCR7 expression is widely considered to be synonymous with thymocyte medullary localization. Based largely on this definition, recent studies indicate that despite the well-established role of the medulla in inducing central tolerance, ∼75% of negative selection occurs in CCR7^-^ ‘cortical’ thymocytes, while only 25% occurs in CCR7^+^ ‘medullary’ cells (Breed et al., 2019; Daley et al., 2013; Hu et al., 2016). It has also been suggested that cortical negative selection may eliminate thymocytes reactive to ubiquitous self-antigens while medullary deletion tolerizes TRA-responsive cells. Our previous research implicates chemokine receptors other than CCR7 in promoting thymocyte medullary localization (Ehrlich et al., 2009). We found that CCR4, which is upregulated by post-positive selection thymocytes, also promotes medullary entry and negative selection (Hu, Lancaster, Sasiponganan, et al., 2015). These findings raise the question of whether CCR4 and CCR7 play distinct or redundant roles in promoting thymocyte medullary entry and central tolerance.

In this paper, we use a combination of approaches, including chemotaxis assays, 2-photon live-cell microscopy, and synchronized positive selection assays, to distinguish the contributions of CCR4 versus CCR7 to thymocyte medullary entry and negative selection. We find that CCR4 is upregulated by thymocytes as early as a few hours after positive selection, while CCR7 is expressed days later by mature post-positive selection cells. CCR4 expression promotes chemotaxis and medullary entry of early post-positive selection thymocytes, which do not yet express CCR7. Initial CCR7 expression results in only moderate thymocyte chemotaxis towards CCR7 ligands and modest accumulation in the medulla, such that these thymocytes migrate in both the cortex and medulla. These findings indicate that CCR7 expression is not a definitive marker of medullary versus cortical localization. CCR7 is, however, required for robust accumulation of mature (CD4SP) thymocytes in the medulla. Notably, consistent with their differential activity in distinct thymocyte subsets, CCR4 and CCR7 are required in early and late phases of polyclonal negative selection, respectively. While CCR7 is known to promote tolerance to TRAs (Kozai et al., 2017; Nitta et al., 2009), we present evidence that CCR4 promotes tolerance to self-antigens presented by activated APCs. Collectively, our study establishes non-redundant roles for CCR4 and CCR7 in governing localization of post-positive selection thymocyte subsets and central tolerance.

## Results

### CCR4 is expressed by immature post-positive selection thymocyte subsets, while CCR7 expression is restricted to more mature thymocyte subsets

We first investigated expression of CCR4 and CCR7 by distinct thymocyte subsets using cell surface markers that delineate developmental stages after positive selection (Figure 1A) (Sinclair et al., 2013; Xing et al., 2016). To validate the developmental trajectory of these thymocyte subsets, we analyzed their GFP expression levels in Rag2p-GFP mice, in which GFP expression is driven by the *Rag2* promoter. In this model, expression of GFP distinguishes newly generated thymocytes (GFP^+^) from recirculating T cells (GFP^-^), and declining GFP expression reflects time elapsed after positive selection, when the *Rag2* promoter becomes inactive (Boursalian et al., 2004). In pre-positive selection CD4^+^CD8^+^ double-positive (DP) CD3^-^CD69^-^ thymocytes, the *Rag2* promoter is active, resulting in maximal GFP expression (Figure 1B). After receiving a TCR signal, thymocytes upregulate CD3 and CD69 (Fu et al., 2009), generating early post-positive selection DP CD3^lo^CD69^+^ cells; the brief time elapsed after positive selection is indicated by continued high GFP expression by this subset (Figure 1B). To further evaluate if DP CD3^lo^CD69^+^ thymocytes represent post-positive selection thymocytes, as opposed to cells undergoing strong TCR signaling driving clonal deletion, we evaluated thymocyte subsets from *Nur77^GFP^* mice, in which GFP levels reflect the strength of TCR signaling (Figure 1—figure supplement 1) (Moran et al., 2011).

**Figure 1.**
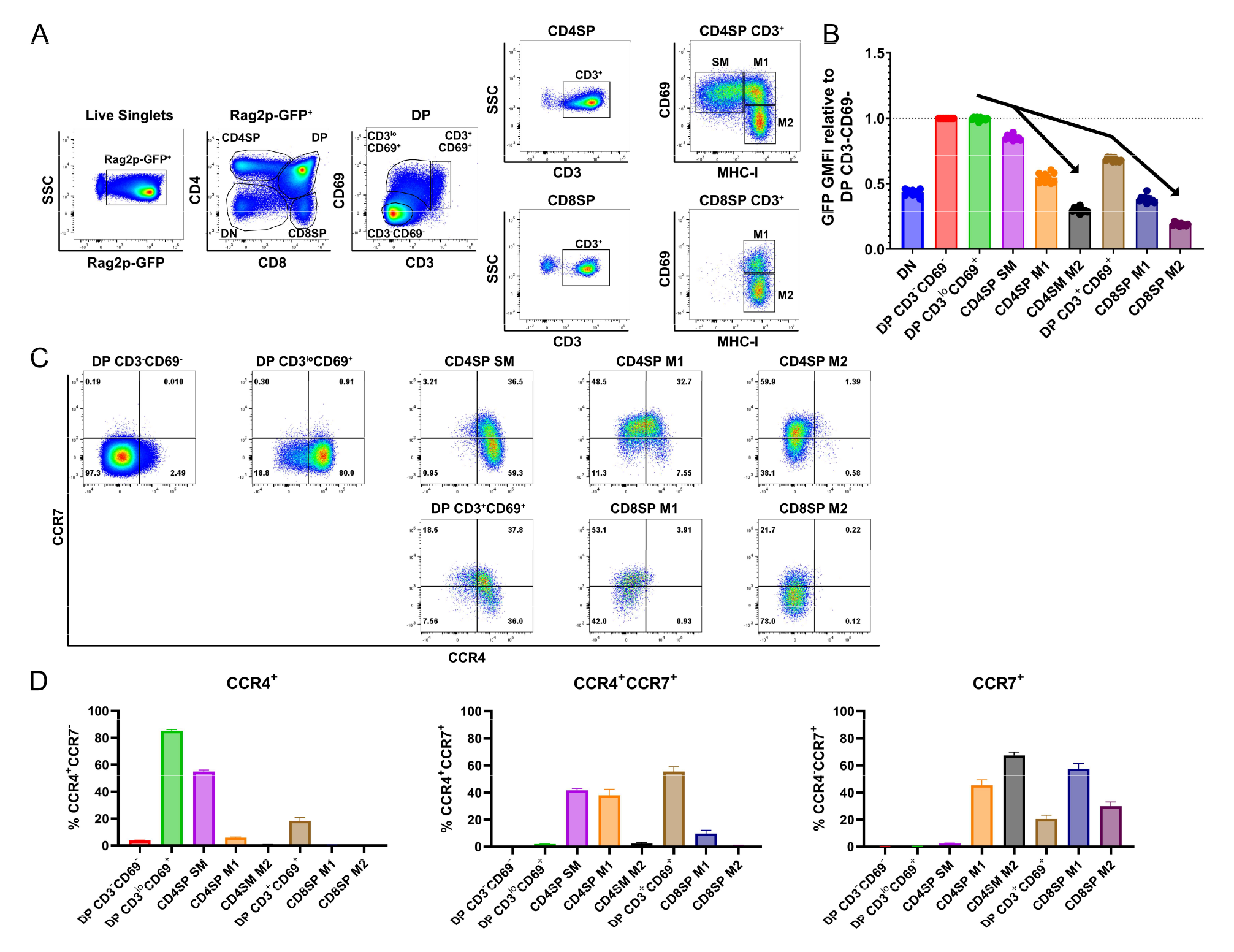
CCR4 and CCR7 are expressed by immature and mature subsets of post-positive selection thymocytes, respectively. **A)** Flow cytometry gating scheme to delineate thymocyte subsets in Rag2p-GFP mice. Post-positive selection DP (CD3^lo^CD69^+^ and CD3^+^CD69^+^), CD4SP (SM, M1, and M2), and CD8SP (M1 and M2) subsets are annotated. **B)** Relative Rag2p-GFP Geometric Mean Fluorescent Intensity (GMFI) of each subset normalized to pre-positive selection DP CD3^-^CD69^-^ cells. Individual data points represent average GFP GMFI of the indicated subset from each mouse. **C)** Representative flow cytometry plots showing CCR4 and CCR7 expression by post-positive selection thymocyte subsets. **D)** Quantification of the percentage of cells of the indicated subset that express CCR4, CCR7, or both chemokine receptors. For B) and D), data were compiled from 4 independent experiments (Mean ± SEM; N=10 mice).

Elevation of GFP expression by DP CD3^lo^CD69^+^ cells, relative to DP CD3^-^CD69^-^ cells, confirmed this subset was TCR-signaled. Thymocytes rescued from negative selection express *Nur77^GFP^* at levels comparable to thymic Tregs, which express self-reactive TCRs (Stritesky et al., 2013). GFP levels were substantially lower in DP CD3^lo^CD69^+^ cells than in CD25^+^ CD4SPs, which are mostly Tregs (Figure 1—figure supplement 1A). Together, these data are consistent with DP CD3^lo^CD69^+^ cells representing mainly post-positive selection DPs, and not strongly self-reactive thymocytes undergoing negative selection. Sequential maturation through by CD4^+^ single-positive semi-mature cells (CD4SP SM) and then CD4SP or CD8SP mature 1 (M1) and mature 2 (M2) subsets was confirmed by progressive reductions in GFP levels for both the CD4 and CD8 lineages (Figure 1B) (Xing et al., 2016). A distinct population of DP CD3^+^CD69^+^ cells expressed lower levels of GFP than CD4SP SM cells (Figure 1B), placing it temporally between CD4SP SM and CD8SP M1 cells (Figure 1B). Thus, we considered whether DP CD3^+^CD69^+^ cells represent MHCI-restricted thymocytes in the process of differentiating into CD8SP cells through co-receptor reversal downstream of the CD4SP SM stage (Brugnera et al., 2000). Consistent with this possibility, DP CD3^+^CD69^+^ cells were significantly reduced in *β2m^-/-^* mice, in which positive selection of CD8SP thymocytes is abrogated, as is selection of innate iNKT cells and CD8αα+ IELps (Bendelac et al., 1994; Ruscher et al., 2017). Furthermore, DP CD3^+^CD69^+^ cells were enriched in *MHCII^-/-^*mice in which only MHCI-restricted thymocytes are positively selected (Figure 1—figure supplement 1B-C)). Together, these data demonstrate that DP CD3^+^CD69^+^ cells are MHC-I restricted and have persisted for an intermediate time between CD4SP SM and CD8SP M1 cells after positive selection, consistent with a co-receptor reversing DP subset.

Having delineated the temporal sequence of thymocyte development post-positive selection, we quantified expression of CCR4 and CCR7 by each subset. CCR4 is upregulated by the majority of early post-positive selection DP CD3^lo^CD69^+^ thymocytes, which do not yet express CCR7 (Figure 1C-D, Figure 1—figure supplement 2A-B). CCR7 is subsequently expressed at high levels by ∼40% CD4SP SM cells, and CCR4 expression persists on this subset (Figure 1C-D). In comparison to *Ccr7*^-/-^ cells, low-level expression of CCR7 can be detected on additional *Ccr7*^+/+^ CD4SP SM cells, consistent with initial upregulation of CCR7 by this subset (Figure 1—figure supplement 2A-B). Expression of CCR4 and CCR7 by MHCI-restricted DP CD3^+^CD69^+^ cells is similar to that of CD4SP SM cells, with slightly more CCR7 and less CCR4 (Figure 1C-D, Figure 1—figure supplement 2A-B). As thymocytes mature through CD4SP and CD8SP M1 and M2 stages, CCR4 is progressively downregulated. CCR7 expression peaks in both CD4SP and CD8SP M1 subsets and is diminished in M2 cells (Figure 1C-D, Figure 1—figure supplement 2A-B). Because cell-surface expression of CCR7 on mature CD4^+^ T cells can be downregulated due to ligand-induced internalization of the chemokine receptor (Britschgi et al., 2008), we tested whether receptor internalization could have obscured detection of CCR7 or CCR4 on ex vivo thymocyte subsets.

Thymocytes were incubated at 37°C for up to 2 hours, to allow re-expression of internalized receptors, before immunostaining with anti-CCR7 and anti-CCR4 antibodies. Increased expression of cell-surface CCR7 or CCR4 was not detected with increasing incubation time (Figure 1—figure supplement 2C-D). These findings confirm that CCR4 expression is an early indicator of positive selection at the DP stage, while CCR7 expression is upregulated later by more mature CD4SP and CD8SP subsets.

### Early post-positive selection thymocytes undergo chemotaxis towards CCR4 ligands, while more mature subsets respond to CCR7 ligands

We next determined whether expression of CCR4 and CCR7 corresponds to thymocyte chemotactic responses to their respective ligands. DP CD3^lo^CD69^+^ cells, which express CCR4 but not CCR7, underwent chemotaxis in response to the CCR4 ligand CCL22, but not the CCR7 ligands CCL19 or CCL21 (Figure 2A- D, Figure 2—figure supplement 1A). CCL22 induced chemotaxis of all subsequent post-positive selection thymocytes, except the most mature M2 thymocyte subsets (Figure 2B, Figure 2—figure supplement 1A), largely in keeping with CCR4 expression patterns. Although CD4SP SM and DP CD3+CD69+ subsets expressed CCR7 (Figure 1C), CCR7-mediated chemotaxis of these subsets did not reach statistical significance when compared to all thymocyte subsets (Figure 2C-D). However, when considered separately, moderate CCR7-mediated chemotaxis of these subsets was revealed, although DP CD3^lo^CD69^+^ cells still did not undergo CCR7-mediated chemotaxis (Figure 2—figure supplement 1A). Consistent with expression of CCR7 by more mature thymocyte subsets, the CCR7 ligands CCL19 and CCL21 induced highly efficient chemotaxis of mature CD4SP and CD8SP M1 and M2 subsets (Figure 2C- D).

**Figure 2.**
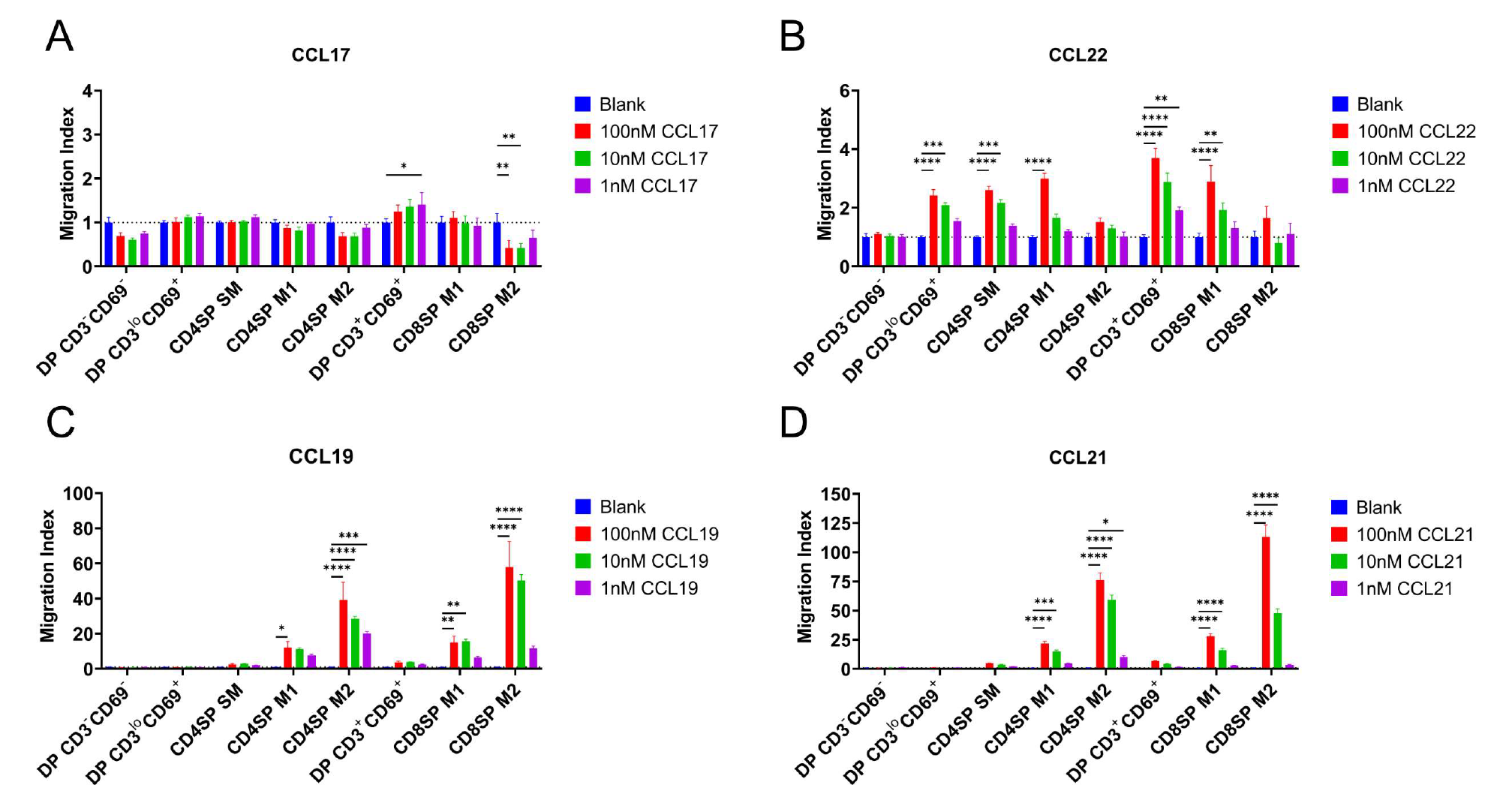
CCR4 and CCR7 promote chemotaxis of immature and mature post-positive selection thymocyte subsets, respectively. **A-D)** Transwell assays were used to quantify chemotaxis of thymocyte subsets to the indicated concentrations of the CCR4 ligands CCL17 (A) and CCL22 (B) and the CCR7 ligands CCL19 (C) and CCL21 (D). Migration indexes were calculated as the frequencies of input cells of each subset that migrated towards the chemokine relative to the frequencies of input cells that migrated in the absence of chemokine (Blank). Data were compiled from 3 independent experiments with triplicate wells per assay. 2-way ANOVA with Dunnett’s multiple comparison correction were used for statistical analysis. (Mean ± SEM; *: p<0.05, **: p<0.01, ***: p<0.001, ****: p<0.0001)

A recent study indicated that expression of the chemokine receptor CXCR4 must be extinguished before post-positive selection thymocytes can leave the cortex, where CXCL12 is expressed by cortical thymic epithelial cells, to migrate into the medulla (Kadakia et al., 2019). Thus, we sought to place CXCR4 expression and function into context with CCR4 and CCR7. CXCR4 expression decreased after positive selection, but all post-positive selection subsets continued to express CXCR4 (Figure 3A-B), and all except for CD4SP SM and DP CD3^+^CD69^+^ subsets underwent chemotaxis in response to CXCL12 (Figure 3C). Although the earliest post-positive selection thymocyte subsets exhibited reduced CXCR4 responsiveness, perhaps allowing them to exit the cortex, the ability to respond to CXCR4 signals does not preclude medullary localization, as evidenced by CXCR4-mediated chemotaxis of medullary CD4SP and CD8SP M2 cells. Taken together, chemotaxis assays show that early post-positive selection DP CD3^lo^CD69^+^ thymocytes are responsive to CCR4 but not CCR7 ligands, with low responsiveness to CXCR4 ligands. At the next stage, CD4SP SM cells also undergo chemotaxis towards CCR4 ligands, and begin to migrate at a low level towards CCR7 ligands, but not to CXCR4 ligands. As thymocytes mature to the CD4SP and CD8SP M1 and M2 stages, they progressively gain the ability to respond robustly to CCR7 ligands, regain CXCR4 responsiveness, and lose responsiveness to CCR4 ligands (Figure 3D).

**Figure 3.**
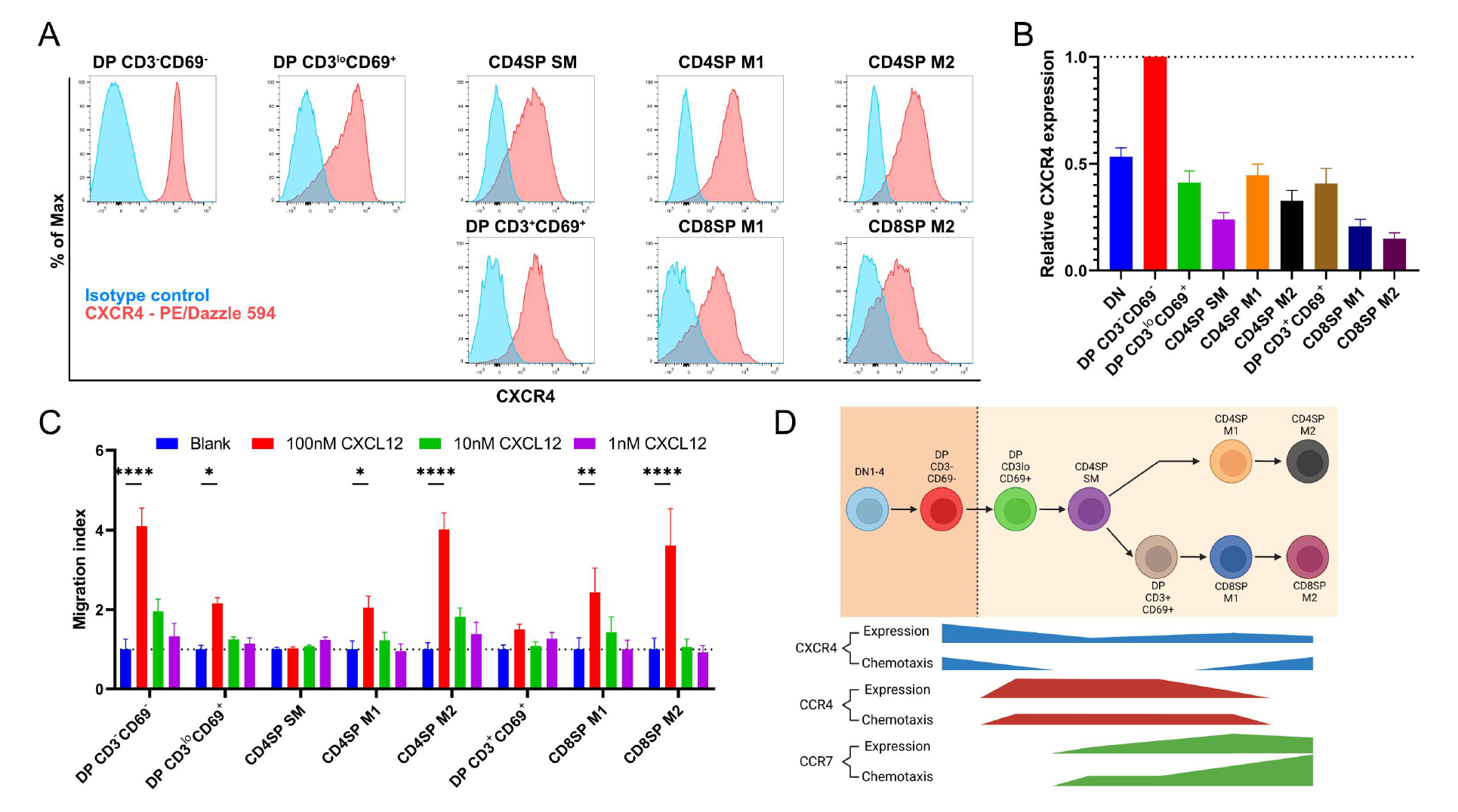
Persistent CXCR4 expression was detected on post-positive selection thymocytes, and CXCR4 activity declines only in intermediate subsets. **A)** Representative flow cytometry plots showing CXCR4 surface expression by the indicated thymocyte subsets, compared to isotype control stains. **B)** Quantification of relative GMFI of CXCR4 for the indicated thymocyte subsets. (Mean ± SEM; N=10) **C)** Transwell assays were used to quantify chemotaxis of thymocyte subsets to the indicated concentrations of CXCL12. Migration indexes were calculated as the frequencies of input cells of each subset that migrated towards the chemokine relative to the frequencies of input cells that migrated in the absence of chemokine (Blank). Data were compiled from 3 independent experiments with triplicate wells per assay. 2-way ANOVA with Dunnett’s multiple comparison correction were used for statistical analysis. (Mean ± SEM; *: p<0.05, **: p<0.01, ***: p<0.001, ****: p<0.0001) **D)** Graphical summary showing relative expression and chemotactic activity of CXCR4, CCR4 and CCR7 on post-positive selection thymocyte subsets based on data in Figures 1-3.

Initial expression of CCR7 by CD4SP SM thymocytes does not result in strong chemotaxis to CCR7 ligands (Figure 2D), and expression of CXCR4 by this subset does not enable detectable chemotaxis towards CXCL12 (Figure 3C), highlighting a temporal disconnect between chemokine receptor expression and function on maturing post-positive selection thymocytes. We considered the possibility that CD4SP SM cells could be intrinsically unable to respond to chemokine receptor signaling. However, these cells responded efficiently to CCR4 ligands (Figure 2B), and they underwent chemotaxis to the CCR9 ligand CCL25 at levels comparable to DP CD3^lo^CD69^+^ cells (Figure 2—figure supplement 2A), consistent with a previous study showing CCL25 induces migration of most thymocyte subsets (Campbell et al., 1999). Thus, regulation of thymocyte responsiveness to chemokine receptors is more complex than altered cell-surface expression levels.

CCL21a establishes a chemokine gradient that drives accumulation of CD4SP thymocytes in the medulla (Kozai et al., 2017; Ueno et al., 2002; Ueno et al., 2004). Because we found that functional CCR4 is expressed by early post-positive selection DP thymocytes, we examined whether CCL22 could establish a similar chemokine gradient to recruit these CCR4-responsive cells into the medulla.

Immunofluorescence analysis revealed that CCL22 is expressed predominantly in the medulla (Figure 2—figure supplement 2B), consistent with a potential role in inducing medullary entry of early post-positive selection thymocytes.

### The timing of CCR4 upregulation following positive selection correlates with thymocyte medullary entry

Given that the majority of early post-positive selection DP thymocytes respond to CCR4 but not CCR7 ligands, we sought to determine how rapidly CCR4 versus CCR7 are upregulated following initiation of positive selection and to assess whether expression of either receptor correlates with the timing of medullary entry. To accomplish these goals, we used a synchronized positive selection thymic slice assay, coupled with flow cytometry or 2-photon microscopy (Figure 4A). OT-II^+^ *Rag2^-/-^ MHCII^-/-^* mice served as a source of pre-positive selection DP thymocytes that did not yet express CCR4 or CCR7 due to lack of positive selection in the absence of MHCII expression (Figure 4B). OT-II^+^*Rag2^-/-^ MHCII^-/-^* thymocytes were fluorescently labeled prior to culturing on MHC-II-sufficient, positively selecting (wild-type (WT or pCX-EGFP) or non-positively selecting (*MHCII^-/-^*) live thymic slices. At various time points, expression of CCR4 and CCR7 by OT-II thymocytes in the thymic slices was assayed by flow cytometry, or medullary entry was quantified by 2-photon live-cell microscopy (Figure 4A).

**Figure 4.**
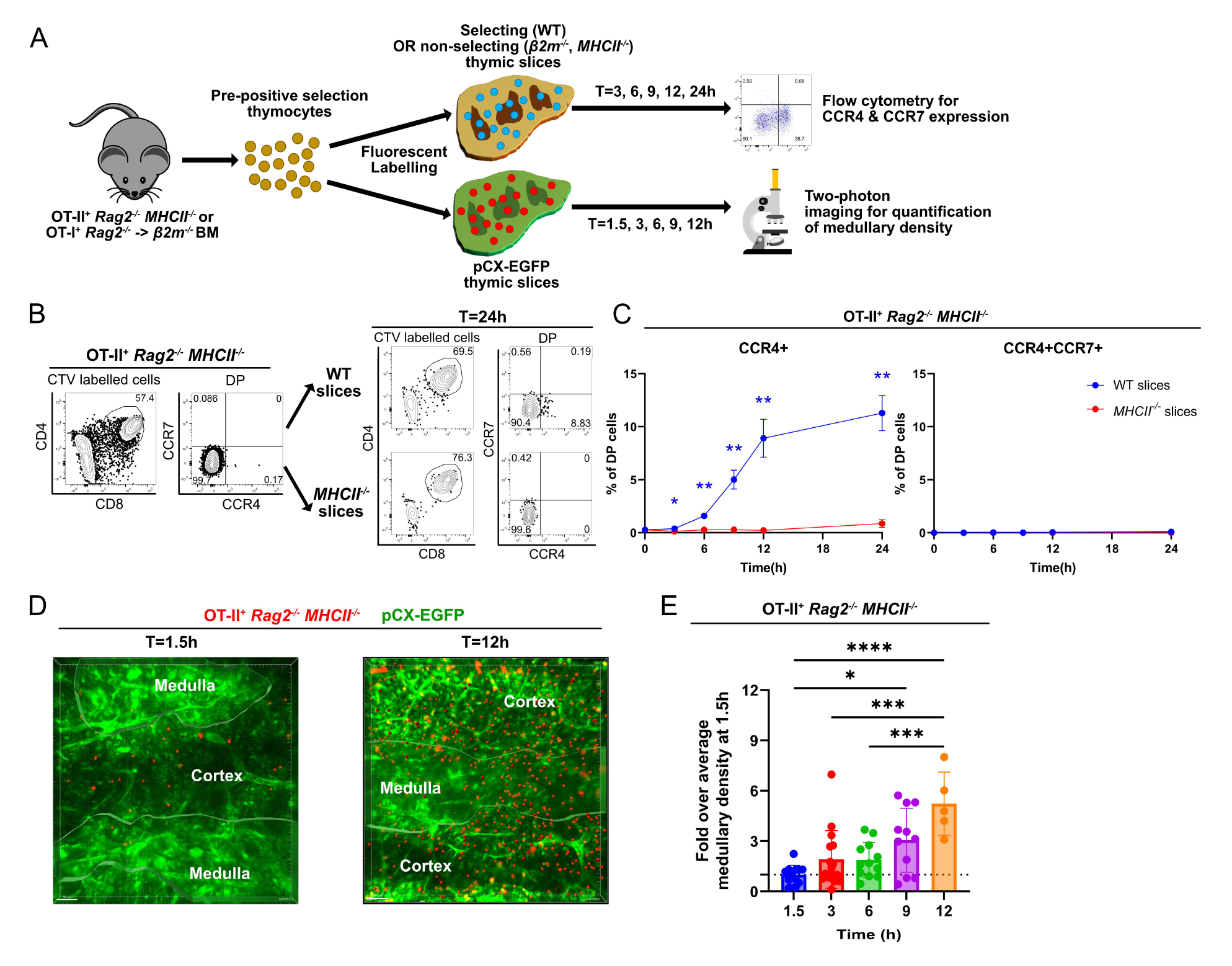
Rapid upregulation of CCR4 following positive selection of OT-II thymocytes correlates with medullary entry. **A)** Experimental schematic of synchronized thymic slice positive selection assay to determine timing of CCR4 and CCR7 expression by flow cytometry and medullary entry by 2-photon microscopy. **B)** Representative flow cytometry plots showing CCR4 and CCR7 expression of OT-II^+^ *Rag2*^-/-^ *MHCII*^-/-^ thymocytes added to thymic slices (left) and analyzed 24h after incubation in WT or *MHCII*^-/-^ slices, as indicated (right). **C)** Percentages of CCR4^+^ and CCR4^+^CCR7^+^ OT-II^+^ thymocytes in thymic slices of the indicated genotypes at the indicated time points. Data were compiled from 3 experiments with triplicate slices each. Mixed-effect analysis with Šídák’s multiple comparison correction was used for statistical analysis. (Mean ± SEM; *: p<0.05, **: p<0.01) **D)** Representative maximum intensity projections of 2- photon imaging data showing a pCX-EGFP thymic slice (green) containing CMTPX-labeled OT-II^+^ *Rag2^-/-^ MHCII*^-/-^ thymocytes (red) at 1.5h and 12h after thymocytes were added to the slices. Medullary and cortical volumes were demarcated as indicated by the masked regions. **E)** Quantification of medullary density of OT-II^+^ *Rag2^-/-^ MHCII*^-/-^ thymocytes at indicated time points after incubation on pCX-EGFP thymic slices. Individual data points represent relative medullary input cell densities from each Video compared to the average of medullary densities from all 1.5h Videos. Data were compiled from 4 time-course experiments. One-way ANOVA with Tukey’s multiple comparison correction were used for statistical analysis. (Mean ± SEM; *: p<0.05, **: p<0.01, ***: p<0.001, ****: p<0.0001)

OT-II^+^ *Rag2^-/-^ MHCII^-/-^* DP cells upregulated CCR4 as early as 3 hours (3h) after being introduced onto positively selecting thymic slices (Figure 4C). The frequency of CCR4^+^ cells continued to increase over 24 hours on WT slices. Positive selection was required for initiation of CCR4 expression, as demonstrated by the failure to upregulate CCR4 on non-selecting *MHCII^-/-^* slices. CCR7 was not upregulated within 24h of initiating positive selection (Figure 4C). When labeled OT-II^+^ *Rag2^-/-^ MHCII^-/-^* thymocytes were introduced into positively selecting pCX-EGFP thymic slices and imaged by 2-photon microscopy, they were initially localized to the cortex, consistent with our previous findings that pre-positive selection thymocytes cannot access the medulla (Video S1) (Ehrlich et al., 2009). By 9-12 hours after positive selection, OT-II^+^ thymocytes entered the medulla, as evidenced by a significant increase in medullary density (Figure 4D- E; Video S2). As CCR7 is not yet expressed by these cells (Figure 4C), CCR4 likely drives medullary entry of early post-positive selection thymocytes.

To test if the timing of CCR4 and CCR7 upregulation and medullary entry are comparable for MHCI-restricted thymocytes, we carried out similar experiments using pre-positive selection OT-I^+^ *Rag2^-/-^* thymocytes that matured in the non-selecting thymuses of β*2m^-/-^* hosts. Although OT-I positive selection was largely inhibited in these bone marrow chimeras, about 10% of the DP cells expressed CCR4 at baseline (Figure 4—figure supplement 1A-B). Nonetheless, CCR4 was upregulated as early as 3h after initiating positive selection in WT thymic slices, but not in non-selecting β*2m^-/-^* slices and increased over 24 hours (Figure 4—figure supplement 1A-B). CCR7 was not expressed during the first 24h after initiation of positive selection and only became detectable as CD8SP cells differentiated at the 72h timepoint (Figure 4—figure supplement 1 B-C), consistent with prior observations (Lutes et al., 2021). Furthermore, 2-photon imaging of pre-selection OT-I^+^ *Rag2^-/-^* thymocytes within the cortex and medulla of thymic slices showed that despite medullary entry of some thymocytes at 1.5h (Video S3), likely due to the observed baseline CCR4 expression in this model, the medullary density of OT-I thymocytes increased significantly by 12h after the initiation of positive selection (Figure 4—figure supplement 1 D-E; Video S4).

Together, these findings show that both MHCI-and MHCII-restricted thymocytes upregulate CCR4 within a few hours of the initiation of positive selection and accumulate significantly in the medulla by 9- 12h. Thymocyte medullary entry precedes CCR7 upregulation, which does not occur until 48h-72h after positive selection. The kinetics of CCR4 upregulation are consistent with a role for CCR4 in promoting early medullary entry of positively selected thymocytes. However, CCR7 is known to be required for efficient medullary entry of SP thymocytes (Ehrlich et al., 2009; Kurobe et al., 2006), raising the possibility that these two chemokine receptors promote medullary localization of different thymocyte subsets.

### CCR4 and CCR7 are required for medullary accumulation of distinct post-positive selection thymocyte subsets

To test our hypothesis that CCR4 and CCR7 are required for medullary accumulation of distinct post-positive selection thymocytes, we FACS purified thymocyte subsets from WT, *Ccr4^-/-^*, *Ccr7^-/-^* and *Ccr4^-/-^ Ccr7^-/-^* (DKO) mice, labeled them with red or blue fluorescent dyes, and allowed them to migrate in pCX-EGFP live thymic slices. 2-photon time-lapse microscopy was used to image the sorted cells, so we could determine their migratory properties and densities in the medulla versus cortex (Figure 5—figure supplement 1A). EGFP is expressed ubiquitously in pCX-EGFP mice, such that the cortex and medulla of thymic slices can be distinguished by differences in stromal cell morphology and EGFP intensity (Figure 5A) (Ehrlich et al., 2009).

**Figure 5.**
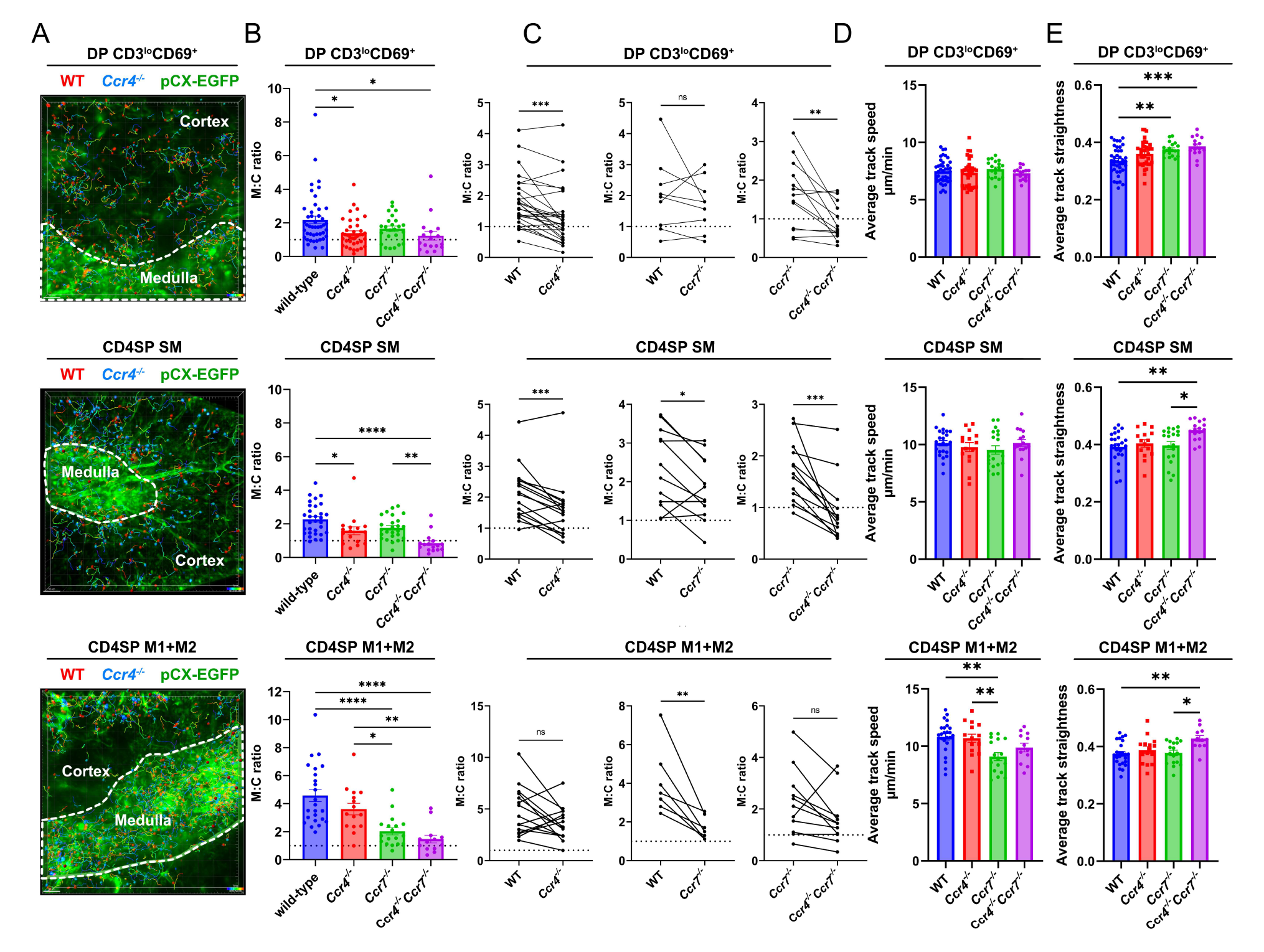
CCR4 and CCR7 are required for medullary accumulation of distinct post-positive selection thymocyte subsets. **A)** Representative maximum intensity projections of 2-photon imaging data showing pCX-EGFP thymic slices (green) containing CMTPX (red)- and Indo-1 AM (blue)-labeled FACS sorted DP CD3^lo^CD69^+^, CD4SP SM or CD4SP M1+M2 cells from WT or *Ccr4^-/-^* mice, as indicated, imaged 1-4 hours after slice entry. The cortex and medulla are demarcated by white dotted lines, and cell tracks are color encoded for elapsed imaging time. **B)** Medullary-to-cortical thymocyte density ratios were quantified from 2-photon imaging data as in A), with individual data points representing ratios from individual Videos. **C)** Paired analysis of medullary-to-cortical density ratios of WT versus *Ccr4^-/-^*, WT versus *Ccr7^-/-^*, or *Ccr7^-/-^* versus *Ccr4^-/-^Ccr7^-/-^* thymocyte subsets imaged simultaneously within the same slices, with individual data points representing ratios from single videos. Paired datasets are taken from the same data shown in B), and paired t-tests were used for statistical analysis. (**: p<0.01, ***: p<0.001) **D-E)** Quantification of D) average thymocyte track speeds and E) average track straightness from imaging data reported in B). Individual data points represent average track speeds or straightness of thymocytes of the indicated genotype in individual Videos. For B), D) and E), data were compiled from 19 independent experiments. One-way ANOVA with Tukey’s multiple comparison correction were used for statistical analysis. (Mean ± SEM; *: p<0.05, **: p<0.01, ***: p<0.001, ****: p<0.0001)

Purified WT DP CD3^lo^CD69^+^ cells entered and migrated within the medulla, accumulating at a 2-fold higher density than in the cortex (Figure 5A-B; Video S5). As expected, the majority of the sorted early post-positive selection cells expressed CCR4, but not CCR7 (Figure 5—figure supplement 1B-C). Medullary enrichment of this subset was abolished when the cells were purified from *Ccr4^-/-^* or DKO mice, but *Ccr7* deficiency did not diminish their medullary accumulation (Figure 5B). Paired analyses of WT versus *Ccr4^-/-^* and *Ccr7*^-/-^ versus DKO DP CD3^lo^CD69^+^ cells imaged together in the same slices confirmed that *Ccr4* deficiency significantly impaired medullary accumulation of this early post-positive selection DP subset irrespective of *Ccr7* genotype, but genetic deficiency of *Ccr7* at this stage did not alter medullary accumulation (Figure 5C). Altogether, CCR4 is required for the 2-fold accumulation of DP CD3^lo^CD69^+^ cells in the medulla relative to the cortex.

Despite the fact that roughly half of the purified CD4SP SM cells expressed CCR7 (Figure 5—figure supplement 1B-C), which is considered to be a hallmark of thymocyte medullary entry, accumulation of these cells within the medulla did not further increase, but instead remained at a 2-fold medullary: cortical density ratio, comparable to that of DPCD3^lo^CD69^+^ cells (Figure 5A-B; Video S6). Medullary accumulation of this subset was significantly reduced by CCR4 deficiency, regardless of the *Ccr7* genotype (Figure 5B-C), indicating an important role for CCR4 in promoting medullary localization of CD4SP SM cells. In addition, paired analysis of *Ccr7^-/-^* versus WT CD4SP SM cells imaged together in the same slices showed that CCR7 also contributes to medullary accumulation of CD4SP SM cells (Figure 5C). Together, these data indicate that CCR4 and CCR7 cooperate to promote medullary accumulation of CD4SP SM thymocytes.

CD4SP M1+M2 thymocytes, the majority of which express only CCR7 (Figure 5—figure supplement 1C), accumulated in the medulla to a greater extent than either of the previous subsets, with a ∼4-5-fold medullary:cortical density ratio (Figure 5B-C; Video S7, S8), consistent with the high migration index of these cells to CCR7 ligands (Figure 2C-D). CCR4 deficiency did not significantly impact medullary accumulation of this subset (Figure 5B-C), consistent with the lack of responsiveness of M2 cells to CCR4 ligands (Figure 2A). However, the trend towards decreased medullary enrichment of *Ccr4* deficient cells may reflect the activity of CCR4 in CD4SP M1 cells (Figure 2A). Medullary accumulation of M1+M2 CD4SP thymocytes was highly dependent on CCR7, as demonstrated by the significant decrease in medullary accumulation of both *Ccr7^-/-^* and DKO cells relative to WT or *Ccr4*^-/-^ cells (Figure 5B, Video S7, S8).

We also considered whether CCR4 and CCR7 alter thymocyte migratory properties. Deficiency in *Ccr4* and/or *Ccr7* did not impact the speed of DP CD3^lo^CD69^+^ or CD4SP SM cells. However, CCR7 deficiency resulted in a significant decline in the speed of CD4SP M1+M2 cells, consistent with our previous finding that CCR7 promotes rapid motility of CD4SP thymocytes (Ehrlich et al., 2009). Interestingly, double deficiency of *Ccr4* and *Ccr7* resulted in a significant increase in the average path straightness of all imaged thymocyte subsets (Figure 5E), suggesting that thymocytes responding to CCR4 and CCR7 ligands migrate in a more tortuous path.

Overall, CCR4 and CCR7 were required for medullary accumulation of distinct post-positive selection thymocyte subsets, largely consistent with their expression patterns and chemotactic function (Figure 3D). Notably, our results show that CCR4 directs early post-positive selection thymocytes into the medulla, demonstrating that lack of CCR7 expression by post-positive selection thymocytes does not preclude medullary localization. Conversely, expression of CCR7 by CD4SP SM cells does not result in enhanced medullary accumulation over the previous subset. Thus, CCR7 expression on its own is not indicative of robust medullary accumulation. Interestingly, only mature CD4SP M1+M2 cells accumulate robustly in the medulla and migrate rapidly in a CCR7-dependent manner.

### CCR4 and CCR7 contribute to early versus late phases of negative selection, respectively

Given the differential impact of CCR4 versus CCR7 on migration and medullary accumulation of post-positive selection thymocyte subsets, we hypothesized that these chemokine receptors are required for negative selection of early versus late post-positive selection cells, respectively. Consistent with our prior observations (Hu, Lancaster, Sasiponganan, et al., 2015), the frequencies of post-positive selection thymocyte subsets were comparable between littermate control WT and *Ccr4^-/-^* mice (Figure 6—figure supplement 1A). CD5 expression levels, which are a proxy for self-reactivity (Azzam et al., 1998; Hawiger et al., 2004; Persaud et al., 2014), were also comparable in the absence of *Ccr4* (Figure 6—figure supplement 1). To test the impact of CCR4 and CCR7 on polyclonal negative selection, we quantified the frequency of cleaved-caspase 3^+^ thymocytes that had undergone TCR signaling in WT, *Ccr4*^-/-,^ *Ccr7*^-/-^, and DKO mice. Intracellular cleaved-caspase 3 in TCR-signaled thymocytes has been shown to mainly reflect negative selection (Breed et al., 2019; Hu et al., 2016), although other apoptotic stimuli, such as glucocorticoid hormones, could also induce cleaved-caspase 3 in thymocytes (Alam et al., 1997; Marchetti et al., 2003). Previous studies showed that negative selection occurs in 2 main phases, generally defined to occur at the CCR7^-^ DP stage in the cortex versus the CCR7^+^ CD4SP stage in the medulla (Breed et al., 2019; Daley et al., 2013; Hu et al., 2016; Stritesky et al., 2013). Thus, we quantified the impact of *Ccr4* versus *Ccr7* deficiency on “early-phase” and “late-phase” negative selection (Figure 6A-B). Interestingly, the majority of early-phase DP cells undergoing negative selection expressed only CCR4, as did ∼60% of late-phase CD4SP cells; thus, the majority of thymocytes undergoing clonal deletion express CCR4, but not CCR7. CCR7 was expressed by ∼40% CD4SP and 60% of CD8SP cells undergoing negative selection (Figure 6B). The frequency of total thymocytes undergoing negative selection declined significantly only in DKO mice (Figure 6C). However, CCR4 deficiency resulted in a significant decrease in the frequency of thymocytes undergoing early-phase, but not late-phase negative selection (Figure 6C). In contrast, CCR7 deficiency resulted in a significantly lower frequency of thymocytes undergoing late-phase negative selection, with no impact on early-phase negative selection. Double-deficiency for CCR4 and CCR7 significantly diminished both early-phase and late-phase negative selection (Figure 6C). It was reported that most negative selection occurs at the earlier phase (Breed et al., 2019; Stritesky et al., 2013), which is consistent with the 2:1 ratio of WT thymocytes undergoing early-to late-phase negative selection (Figure 6D). This ratio falls significantly in *Ccr4*^-/-^ mice and increases significantly in *Ccr7*^-/-^ and DKO mice, highlighting the respective contributions of CCR4 and CCR7 to early versus late phases of negative selection, respectively (Figure 6D). Altogether, these data indicate that, consistent with its role in promoting medullary entry of early-post positive selection thymocytes, CCR4 is required for efficient early-phase negative selection of polyclonal DP cells. In contrast, CCR7, which promotes medullary accumulation of more mature CD4SP and CD8SP cells, is required for efficient negative selection of late-phase CD4SP and CD8SP cells.

**Figure 6.**
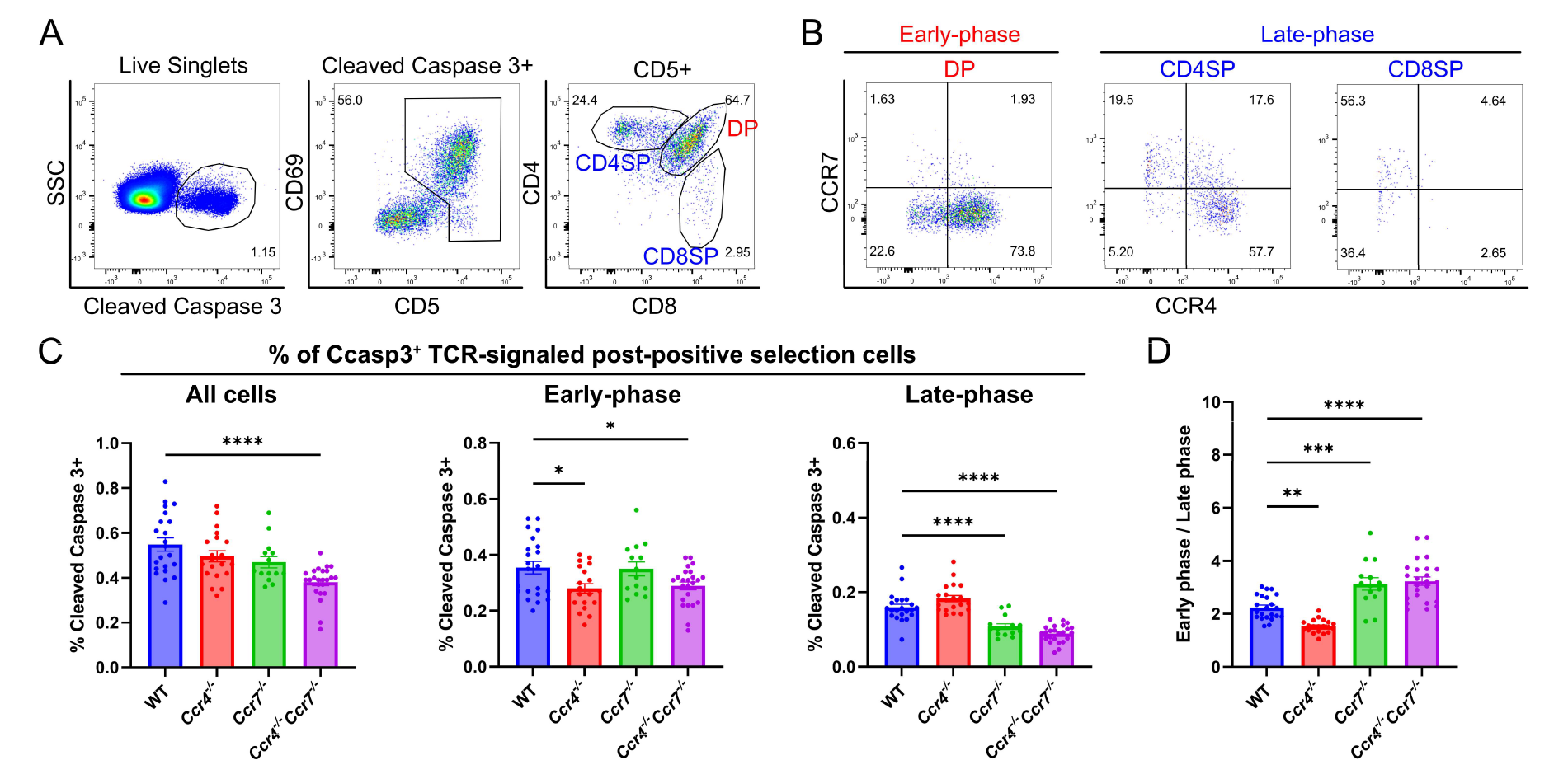
CCR4 and CCR7 contribute to early versus late phases of negative selection, respectively. **A)** Flow cytometric gating scheme to quantify thymocytes undergoing early-phase (DP) or late-phase (CD4SP+CD8SP) negative selection. Representative data from a WT mouse is shown. **B)** Representative flow cytometry plots showing CCR4 and CCR7 expression by thymocytes undergoing early-and late-phase negative selection, as gated in C). **C)** Frequencies of total, early-phase (DP), and late-phase (CD4SP+CD8SP) negative selection among all live thymocytes in WT, *Ccr4^-/-^*, *Ccr7^-/-^*, and *Ccr4^-/-^Ccr7^-/-^* mice. **D)** Ratio of thymocytes undergoing early-to late-phase negative selection in WT, *Ccr4^-/-^*, *Ccr7^-/-^*, and *Ccr4^-/-^Ccr7^-/-^* mice. For C) and D), One-way ANOVA with Dunnett’s multiple comparison correction were used for statistical analysis. (n=22 for WT, n=20 for *Ccr4^-/-^*, n=14 for *Ccr7^-/-^*, n=24 for *Ccr4^-/-^ Ccr7^-/-^*. Mean ± SEM; *: p<0.05, **: p<0.01, ***: p<0.001, ****: p<0.0001)

### CCR4 and CCR7 suppress autoinflammation in distinct tissues, with CCR4 promoting tolerance to activated antigen-presenting cells

Given that CCR4 and CCR7 enforced central tolerance at distinct stages of T-cell development when thymocytes might interact with different APCs, these chemokine receptors could promote tolerance to distinct self-antigens. To evaluate this possibility, we assessed the presence of spontaneous autoimmunity/autoinflammation in multiple organs of WT, *Ccr4^-/-^*, *Ccr7^-/-^* and *Ccr4^-/-^Ccr7^-/-^*mice. We previously found that *Ccr4^-/-^* and *Ccr7^-/-^* mice had anti-nuclear autoantibodies by ∼12-months of age (Hu et al., 20115). Here we detected anti-nuclear autoantibodies in the serum of 67% of *Ccr4^-/-^*, 100% of *Ccr7^-/-^, and 75% of Ccr4^-/-^Ccr7^-/-^ mice* between 5 to 6.5 months of age, relative to 22% of WT mice (Figure 7—figure supplement 1A). Thus, we evaluated other organs from mice in this age range for lymphocytic infiltrates or inflammation. Colon inflammation was not observed in *Ccr7*^-/-^ or WT mice. Colons from *Ccr4*^-/-^ mice showed mild epithelial hyperplasia, with small foci of acute inflammation. In contrast, Ccr4^-/-^ *Ccr7*^-/-^ mice demonstrated moderate to severe colonic mucosal hyperplasia, with abundant acute and chronic inflammation extending into the submucosa. Active ulcerations were observed in 40% of *Ccr4^-^*^/-^ *Ccr7*^-/-^ mice, with crypt abscesses and regenerative changes consistent with prior ulceration in most *Ccr4*^-/-^ *Ccr7*^-/-^ mice (Figure 7A, left). Histologic scores were markedly and significantly higher in *Ccr4^-/-^ Ccr7^-/-^* mice compared to WT, with a trend toward higher scores in *Ccr4^-/-^* mice that is absent in *Ccr7^-/-^* mice (Figure 7A, right), suggesting that CCR4 and CCR7 collaboratively maintain self-tolerance in the colon (Figure 7A). Immune infiltrates were elevated in the liver, lacrimal glands, and submandibular glands of *Ccr7^-/-^* and *Ccr4^-/-^Ccr7^-/-^* mice, but not *Ccr4^-/-^* mice at 6-months of age (Figure 7B, Figure 7- figure supplement 1B-D), indicating that CCR7 plays a dominant role in promoting tolerance to endocrine organs, consistent with the established requirement for CCR7 to mediate negative selection against mTEC-expressed TRAs, but not ubiquitous self-antigens (Nitta et al., 2009). Interestingly, 70-75% of *Ccr4*^-/-^ mice, compared to 17-20% of WT and 0% of *Ccr7*^-/-^ mice displayed moderate lymphoid hyperplasia in the spleen and mesenteric lymph nodes of 12-month old mice (Figure 7C, right). Immunofluorescent staining of lymph nodes from 6-month old *Ccr4*^-/-^ mice revealed abnormal CD4 and CD8 distributions/expansions in T-cell zones relative to WT mice (Figure 7C, left). As expected, T cell zones from *Ccr7^-/-^* mice were hypoplastic (not shown), reflecting the role of CCR7 in mediating T-cell entry into LNs. These results indicate that CCR4 maintains T cell homeostasis in secondary lymphoid organs.

**Figure 7.**
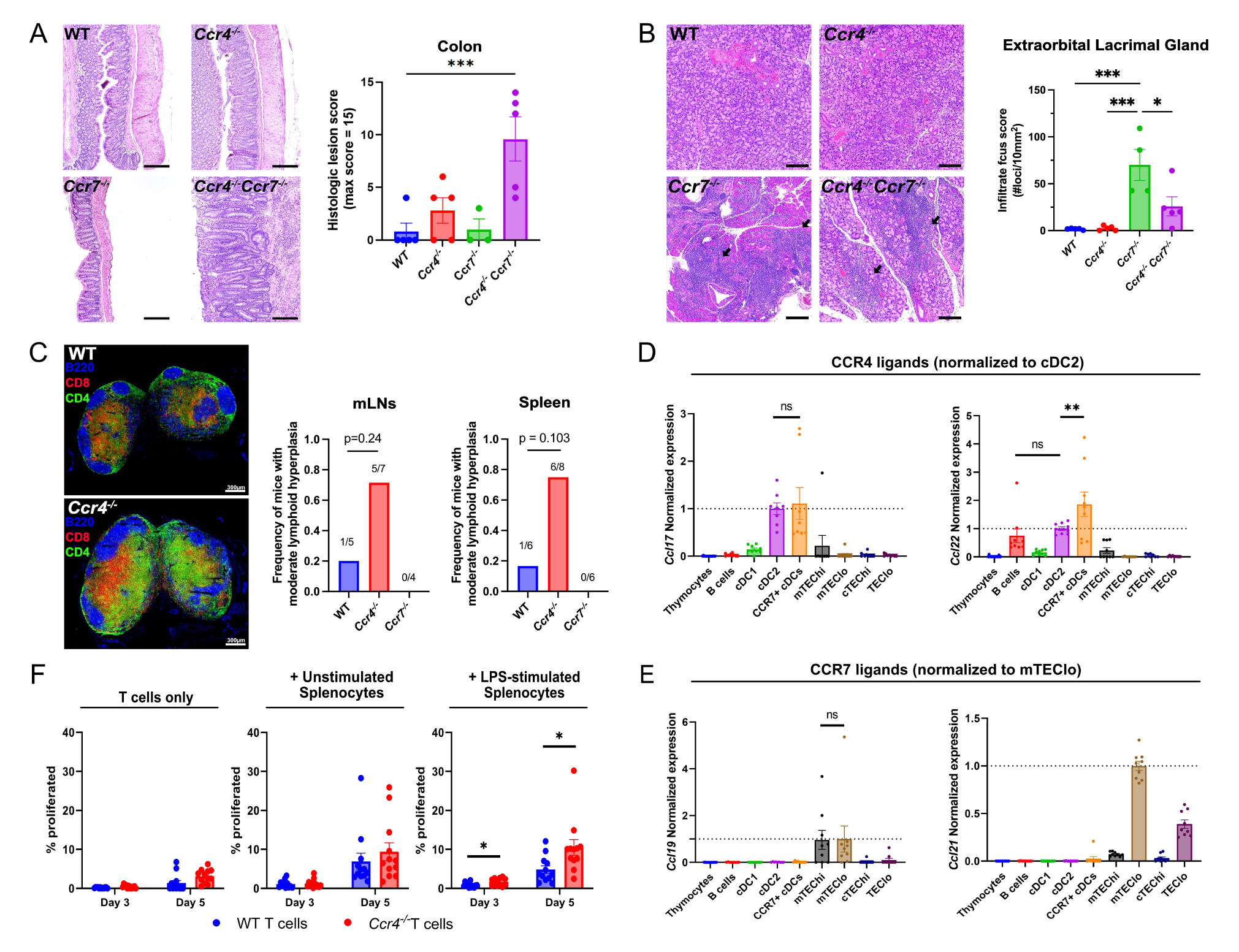
CCR4 and CCR7 promote tolerance to distinct organs, with CCR4 promoting tolerance to activated antigen-presenting cells. **A)** Representative images of H&E stains from the colon of 5–6-month-old WT, *Ccr4^-/-^*, *Ccr7^-/-^* or *Ccr4^-/-^ Ccr7^-/-^* mice (left). Bars, 200 µm. Histological lesion score was quantified by a pathologist blinded to treatment group (right). One-way ANOVA was used to determine significance between groups (Mean ± SEM; ***: p<0.001).**B)** Representative images of H&E stains from extraorbital lacrimal glands of 5–6- month-old WT, *Ccr4^-/-^*, *Ccr7^-/-^* or *Ccr4^-/-^Ccr7^-/-^* mice (left). Bars, 500 µm. Arrows indicate lymphocytic infiltrate foci. The number of infiltrate foci per 10mm^2^ was quantified (right), and one-way ANOVA was used to test significance between groups (Mean ± SEM; *: p<0.05, ***: p>0.001): C) Representative confocal images of CD4 (green), CD8 (red), and B220 (blue) on immunostained cryosections of inguinal lymph nodes from 5-6 month-old WT and *Ccr4*^-/-^ mice(left). Frequency of moderate lymphoid hyperplasia in mLNs and spleens from 12-13 month-old WT, *Ccr4^-/-^*and *Ccr7^-/-^* mice, as determined by a veterinary pathologist (right). Fisher’s exact test was used to determine significance. **D-E)** Expression of CCR4 ligands (D) and CCR7 ligands (E) by distinct thymic antigen presenting cell subsets, assessed by qRT-PCR. Expression levels were normalized to subsets previously reported to expresses the respective ligands: CCL17 and CCL22 expression were normalized to cDC2 (Hu, Lancaster, Sasiponganan, et al., 2015), and CCL19 and CCL21 expression to mTEClo (Ki et al., 2014; Misslitz et al., 2004). Data were compiled from 3 independent experiments with triplicates. One-way ANOVA with Dunnett’s multiple comparison correction were used for statistical analysis. (Mean ± SEM; ns: p>0.05, **: p<0.01) **F)** The percentage of WT or *Ccr4^-/-^* T cells that proliferated at day 3 and day 5 when cultured without splenocytes, with unstimulated splenocytes, or with LPS-stimulated splenocytes, as indicated. Data were compiled from 4 independent experiments with triplicate wells, and unpaired t-tests with Holm-Šídák’s multiple comparison correction was used for statistical analysis. (Mean ± SEM; *: p<0.05)

Given the distinct contributions of CCR4 and CCR7 to establishment of central tolerance and maintenance of self-tolerance across different organs, we considered the possibility that CCR4 and CCR7 might promote thymocyte selection against antigens presented by different types of thymic APCs. Thymic APC subsets were FACS purified (Figure 7—figure supplement 2A), and expression of CCR4 and CCR7 ligands was quantified by qRT-PCR. Consistent with our previous study (Hu, Lancaster, Sasiponganan, et al., 2015), the CCR4 ligands CCL17 and CCL22 were expressed by Sirpα^+^ cDC2 cells.

Activated thymic CCR7^+^ cDCs (Ardouin et al., 2016; Hu et al., 2017; Oh et al., 2018) also expressed CCR4 ligands, with the highest levels of CCL22 expressed by activated DCs (Figure 7D). Thymic B cells also expressed CCL22, consistent with a recent report that CCL22 expressed by activated B cells promotes interactions with T cells in the germinal center (Liu et al., 2021). The CCR7 ligands CCL19 and CCL21 were expressed by mTEC subsets, with the highest levels of CCL21 expression by mTEC^lo^ cells (Figure 7E), consistent with previous reports (Bornstein et al., 2018; Kozai et al., 2017). These results suggest that CCR4 may promote thymocyte interactions with B cells and cDCs, including activated CCR7^+^ cDCs that express elevated levels of MHCII and co-stimulatory molecules (Ardouin et al., 2016; Hu et al., 2017), while CCR7 may promote thymocyte interactions primarily with mTECs.

Because a large proportion of thymic B cells (Cepeda et al., 2018) and cDCs (Ardouin et al., 2016; Hu et al., 2017) have activated phenotypes, and T-cell zones in the lymph nodes of *Ccr4^-/-^*mice are hyperplastic (Figure 7C), we considered the possibility that CCR4-directed negative selection could facilitate establishment of central tolerance to antigens expressed by activated APCs. To explore this hypothesis, CD45.1^+^ congenic splenocytes were stimulated overnight with toll-like receptor (TLR) ligands. We confirmed that stimulation with TLR ligands induced activation of the splenic cDCs and B cells based on upregulation of MHCII, CD80, and CD86 (Figure 7—figure supplement 2B). The activated splenocytes were then co-cultured with CD45.2^+^ CD4^+^ conventional T cells isolated from WT or *Ccr4^-/-^* spleens, and the frequency of T cells induced to proliferate was quantified after 3-5 days (Figure 7F). Notably, LPS-stimulated splenocytes induced more proliferation of *Ccr4^-/-^* versus WT CD4^+^ T cells at days 3 and 5 (Figure 7G). Splenocytes that were unstimulated or activated by other TLR ligands did not preferentially induce *Ccr4*^-/-^ T cell proliferation (Figure 7E, Figure 7—figure supplement 2C). Taken together, these data suggest that CCR4 promotes thymocyte central tolerance to self-antigens expressed by LPS-activated APCs, possibly by promoting interactions with such APCs in the thymus.

## Discussion

Our data demonstrate that CCR4 and CCR7 play distinct roles in thymocyte localization and the induction of T-cell central tolerance. We find that CCR4 is upregulated by thymocytes within a few hours of the initiation of positive selection, largely coincident with the timing of medullary entry. CCR4 is expressed by almost all early post-positive selection DP CD3^lo^CD69^+^ cells and is required for their 2-fold accumulation in the medulla and efficient negative selection. Notably, the majority of thymocytes undergoing negative selection are DP CD3^lo^CD69^+^ cells, which do not express CCR7. Thus, while the role of CCR4 in negative selection has been questioned (Cowan et al., 2014), our analyses reveal an important role for this chemokine receptor in localization and central tolerance of early post-positive selection polyclonal thymocytes, in keeping with medullary localization of the CCR4 ligand CCL22. In contrast, our study and a previous report (Lutes et al., 2021) show that CCR7 is not upregulated until 48- 72 hours after positive selection. CCR7 is expressed by about half of the CD4SP SM cells, which continue to express CCR4 and respond to CCR4 ligands. In keeping with dual expression, CCR4 and CCR7 cooperate to induce the 2-fold accumulation of CD4SP SM cells in the medulla. As thymocytes mature further to the CD4SP and CD8SP M1 and M2 stages, CCR7 is upregulated, CCR4 is downregulated, and the cells become increasingly chemotactic to CCR7 ligands. We find that CCR7 is required for the robust accumulation of CD4SP M1+M2 cells in the medulla and for late-phase negative selection of CD4SP and CD8SP thymocytes. The finding that the overall frequency of negative selection in the thymus is reduced significantly only in the absence of both CCR4 and CCR7 further indicates that these chemokine receptors both contribute to the induction of central tolerance. Altogether, our findings show a temporally correlated shift after positive selection from a requirement for CCR4 to a requirement for CCR7 for thymocyte medullary localization and negative selection.

Several of our findings raise questions about the current model of negative selection, which assumes that CCR7 expression is a surrogate indicator of thymocyte medullary versus cortical localization (Breed et al., 2019; Hu et al., 2016). This assumption is based on studies, including our own, showing that CCR7 is expressed by CD4SP and CD8SP thymocytes, which are localized mainly within the thymic medulla at steady-state, as evidenced by immunostaining (Ehrlich, 2016). In addition, prior studies showed that accumulation of CD4SP cells in the medulla is CCR7-dependent (Ehrlich et al., 2009; Ueno et al., 2004); consistent with these findings, we confirm that robust medullary accumulation of the more mature CD4SP M1 and M2 cells is CCR7-dependent. Equating CCR7 expression with medullary localization has led to a model in which the early phase of negative selection is thought to impact CCR7^-^ DP cells in the cortex, while the later phase impacts more mature CCR7^+^ cells in the medulla (Breed et al., 2019; Hu et al., 2016). In support of cortical negative selection, apoptotic TCR transgenic cells were located in the cortex of the H-Y^CD4^ model of negative selection (McCaughtry et al., 2008). Also, TCR-signaled thymocytes are present at the cortical side of the CMJ in *Bim*^-/-^ mice, in which apoptosis following strong TCR signaling is impaired (Stritesky et al., 2013), suggesting thymocytes encounter negatively selecting self-antigens in the cortex. Furthermore, CCR7 deficiency impairs negative selection to TRAs, but not to ubiquitous self-antigens (Kurobe et al., 2006; Nitta et al., 2009), indicating that CCR7^+^ mature thymocytes undergo negative selection in the medulla, where TRAs are expressed by mTECs.

While our data confirm that CCR7 is critical for medullary accumulation of more mature thymocyte subsets, we find that CCR7 expression is not a reliable indicator of medullary localization in earlier post-positive selection thymocytes. CCR7^-^ DP CD3^lo^CD69^+^ enter the medulla, where they accumulate in a CCR4-dependent manner with a 2-fold higher density than in the cortex, so CCR7 is not required for medullary entry. Conversely, although almost half of CD4SP SM cells express CCR7, their medullary: cortical density is also 2-fold. CCR7 deficiency only modestly diminished medullary accumulation of CD4SP SM cells, consistent with their moderate chemotaxis towards CCR7 ligands; instead, CCR4 played a greater role in medullary accumulation of this subset. Thus, the two earliest post-positive selection subsets accumulate in the medulla at similar densities, despite CCR7 upregulation by one of these subsets. DP CD3^lo^CD69^+^ and CD4SP SM cells both migrate in the cortex and in the medulla and could thus encounter APCs presenting cognate self-antigens that drive negative selection in either region. Based on the 2-fold enrichment of DP CD3^lo^CD69^+^ cells in the medulla, a substantial proportion of CCR7^-^ early phase negative selection may occur in the medulla. Moreover, some late phase negative selection, defined by either CCR7 expression or maturation to the CD4SP SM stage, may occur in the cortex. These data indicate that CCR7 expression cannot be used as a reliable surrogate biomarker of medullary localization. The location in which thymocytes undergo negative selection impacts which APC subsets will be encountered, altering the spectrum of self-antigens to which thymocytes are tolerized. Thus, it will be important to determine where distinct thymocyte subsets undergo negative selection and which APCs promote tolerance of these different subsets.

It has recently been suggested that CXCR4 expression must be extinguished after positive selection to release thymocytes from binding to cTECs in the cortex, thus enabling medullary entry (Kadakia et al., 2019). However, we found that while CXCR4 is partially downregulated on the cell surface of post-positive selection thymocytes, all subsets except CD4SP SM and DP CD3^+^ CD69^+^ cells undergo chemotaxis to CXCL12, despite their accumulation in the medulla. These findings indicate that CXCR4 expression and activity are compatible with medullary localization. This seeming discrepancy may reflect the overexpression system used to reveal that persistent CXCR4 expression prevented thymocyte medullary entry, perhaps resulting in superphysiological tethering of cells to the cortex (Kadakia et al., 2019). In keeping with the ability of CXCR4^+^ thymocytes to enter the medulla, higher CXCR4 surface expression and responsiveness are observed in mature CD4SP and CD8SP M1 and M2 subsets, which accumulate in the medulla to the greatest extent. It is notable that chemotactic activity of CXCR4 is completely extinguished at the CD4SP SM stage, despite continued CXCR4 expression, but not at the more mature CD4SP and CD8SP stages, suggesting a thymocyte-intrinsic mechanism of diminished responsiveness to CXCR4, similar to the lack of responsiveness to CCR7 ligands by this subset. It is possible that the lack of CXCR4 activity in CD4SP SM cells may permit medullary entry, consistent with the need to release post-positive selection thymocytes from binding to cTECs (Kadakia et al., 2019); however, DP CD3^lo^CD69^+^ cells enter the medulla just as efficiently as CD4SP SM cells despite continued CXCR4 responsiveness, suggesting factors other than reduced CXCR4 activity likely enable medullary entry. The role for increased CXCR4 activity in more mature SP thymocytes remains to be determined but could impact thymocyte egress. Identifying intrinsic regulators of chemokine receptor responsiveness in different thymocyte subsets will further our understanding of the complex mechanisms underlying the orchestration of thymocyte movement and selection in the thymus.

A major outstanding question raised by our findings is whether the combinatorial and temporally regulated changes in chemokine receptor expression on post-positive selection thymocytes drive interactions with distinct APC subsets. Here we show that CCR7 ligands are expressed by mTECs, while CCR4 ligands are expressed mainly by cDC2s and activated cDCs, with some expression by thymic B cells. Thus, CCR7 activity may drive interactions with mTECs, possibly explaining why CCR7 is particularly required for negative selection to TRAs, consistent with the finding that CCR7 deficiency results in autoimmune infiltrates in endocrine organs, as occurs in Aire-deficient mice (Anderson et al., 2002; Kurobe et al., 2006; Nitta et al., 2009). In contrast, CCR4 activity may promote interactions with and tolerance to cDC2s, B cells, and activated cDCs, consistent with our finding that *Ccr4*^-/-^ mice showed T cell hyperplasia in secondary lymphoid organs and that *Ccr4*^-/-^ T cells undergo increased proliferation to LPS-activated splenic APCs. Recently, a subset of cDC2s has been shown to present circulating antigens to induce negative selection, and another subset has been shown to traffic microbial antigens to the thymus, impacting thymocyte selection (Atibalentja et al., 2009; Vollmann et al., 2021; Zegarra-Ruiz et al., 2021). B cells also present self-antigens to induce thymocyte negative selection (Perera et al., 2016; Yamano et al., 2015). While we find CCL22 is expressed predominantly in the medulla, and CCR4 promotes medullary accumulation of early-post positive selection DPs, our study does not rule out the possibility that CCR4 could also enforce interactions between CCR4-responsive thymocytes and cortical DCs expressing CCL22, which could contribute to cortical negative selection. This possibility is particularly intriguing in light of the fact that DP CD3^lo^CD69^+^ and CD4SP SM cells respond to CCR4 ligands and migrate both in the cortex and medulla. The finding that CCR4 and CCR7 cooperate to prevent inflammation in the colon is unexpected and warrants further exploration. Whether the inflammatory phenotypes observed reflect impaired central tolerance due to diminished interactions between thymocytes and distinct thymic APCs and/or impaired peripheral tolerance remains to be determined.

Altogether, our findings suggest a layered process of central-tolerance induction in which CCR4 first promotes interactions with DCs and B cells, driving self-tolerance to activated APCs, which may be present in both the cortex and/or medulla. As thymocytes mature further, CCR7 is expressed and becomes active, drawing the cells robustly into the medulla, where the partially tolerized repertoire can be focused on tolerance induction to the sparse TRAs. One limitation of our current system is that ex vivo live thymic slices are not equivalent to in situ thymi, as they lack circulation, for example. We note, however, that short-term live thymic slice cultures have been widely used to investigate the development, localization, migration, and positive and negative selection of thymocytes, as they have been shown to faithfully reflect these in vivo processes, including confirming that CCR7 signaling induces chemotaxis of mature thymocytes from the cortex into the medulla (Au-Yeung et al., 2014; Dzhagalov et al., 2013; Ehrlich et al., 2009; Lancaster et al., 2019; Melichar et al., 2013; Ross et al., 2014). Because our study measured differences in medullary accumulation of thymocytes that differed genetically only in expression of CCR4 and/or CCR7, we attribute altered localization to the impact of these chemokine receptors on thymocyte migration. However, we note the possibility that CCR4 and CCR7 could signal in conjunction with survival cues, like cytokines, localized to different thymic microenvironments, which could also impact survival of thymocytes in different regions. As thymocytes were imaged within a few hours of entering slices, differential survival in the cortex and medulla due to cytokine cues is somewhat less likely than differential migration. Multiple aspects of our revised model of thymocyte migration and central tolerance, including the impact of CCR4 and CCR7 on interactions with distinct APCs during tolerance induction and on selection of distinct TCR clones will be tested in future studies.

## Materials and Methods

### Mice

C57BL/6J (wild-type), B6.SJL-Ptprc^a^ PepC^b^ (CD45.1), C57BL/6-Tg(TcraTcrb)1100Mjb/J (OT-I), B6.Cg-Tg(TcraTcrb)425Cbn/J (OT-II), B6(Cg)-Rag2^tm1.1Cgn^/J (*Rag2^-/-^*), B6.129P2(C)Ccr7^tm1Rfo^r/J (*Ccr7^-/-^*), B6.129S2- H2^dlAb1-Ea^/J (*MHCII^-/-^*), B6.129P2-B2m^tm1Unc^/DcrJ (*β2m^-/-^*), C57BL/6-Tg(Nr4a1-EGFP/cre)820Khog/J (Nur77^GFP^) were purchased from the Jackson Laboratory. *Ccr4^-/-^*(Chvatchko et al., 2000), pCX-EGFP (Wright et al., 2001), and Rag2p-GFP (Boursalian et al., 2004) strains were generously provided by A.D. Luster (Massachusetts General Hospital, Boston, MA), Irving L. Weissman (Stanford University, Stanford, CA), and Ellen R. Richie (the University of Texas MD Anderson Center, Houston, Texas, United States), respectively. *Ccr4^-/-^ Ccr7^-/-^*, OT-I *Rag2^-/-^*, and OT-II *Rag2^-/-^ MHCII^-/-^* strains were bred in house. Experiments were performed using mice 4-8 weeks old, except for for autoimmune/inflammatory studies which were carried out in older mice, as specified. All strains were bred and maintained under specific pathogen-free conditions at the Animal Resources Center, the University of Texas at Austin, with procedure approval from the Institutional Animal Care and Use Committee, the University of Texas at Austin.

### Flow cytometry

5×10^6^ thymocytes were stained with fluorescently conjugated antibodies and a viability dye, and incubated on ice for 30 minutes in the dark. Unless specified, for stains including anti-CCR7, cells were instead incubated in a 37°C water bath for 45 minutes in the dark. Stained cells were washed and resuspended in FACS Wash Buffer (PBS + 2% FBS (GemCell™, Gemini, CA, USA)) + 1µg/mL propidium iodine (PI) if a fixable viability dye was not used. For cleaved caspase 3 stains, surface-stained cells were fixed and permeabilized with the BD Cytofix/Cytoperm kit (BD, NJ, USA) according to manufacturer’s instructions prior to staining for anti-cleaved caspase 3 (Cell Signaling Technology, MA, USA). All flow cytometry data were acquired on an LSR Fortessa flow cytometer (BD) and analyzed with FlowJo ver.10.8.0 (BD).

### Transwell Chemotaxis assays

Thymocyte chemotaxis assays were performed as previously described (Campbell et al., 1999). Briefly, 5×10^5^ thymocytes were resuspended in 100µL RPMI 1640 (Gibco, MA, USA) + 10% Fetal Bovine Serum (Gemini) and added to the top chamber of 5µm pore transwell tissue culture inserts (Corning, NY, USA) in 24-well plates. The bottom chamber of the transwells contained recombinant mouse chemokines CCL17, CCL22, CCL19, CCL21 (Peprotech, NJ, USA) or CCL25 (R&D Systems, MN, USA) at 100nM, 10nM and 1nM, diluted in 500µL RPMI 1640 + 10% FBS. The plate was cultured at 37°C, 5% CO_2_ for 2 hours. Cells that migrated to the bottom wells, and 5×10^5^ input cells, were analyzed and quantified by flow cytometry. A standard number of 15µm polyester beads were added to each well to calculate the absolute number of thymocytes that migrated. Migration percentages of each subset were calculated by dividing the number of migrated cells of each subset by the number of cells of the same subset in the input cell sample, and the migration index was calculated as the ratio of the migration percentage in wells containing chemokine to the average migration percentage in wells without chemokine (Blank wells).

### *Ex vivo* thymic slice preparation

*Ex vivo* thymic slices were prepared as previous described (Lancaster & Ehrlich, 2017). Briefly, surgically removed thymi were cleaned of residual connective tissue, and the two lobes were separated and embedded in 4% low melting point agarose (Lonza, NJ, USA). 400µm thymic slices were generated by slicing the trimmed agarose blocks on a VT 1000S vibratome (Leica, Germany) and kept on ice, submerged in complete RPMI until addition of thymocytes. Slices were placed on 0.4µm cell culture inserts (Millipore, MA, USA) in 35mm dishes containing 1mL complete RPMI (RPMI 1640 supplemented with 1X GlutaMAX, 1mM Sodium Pyruvate, 1X Penicillin-Streptomycin-glutamine, 1X MEM Non-Essential Amino Acid, and 50µM β-mercaptoethanol (All from Gibco), and 10% FBS (Gemini)).

### Synchronized positive selection thymic slice assays

Synchronized positive selection assays were set up as previously described (Ross et al., 2014). Briefly, to generate a source of pre-positive selection OT-I thymocytes, bone marrow from OT-I *Rag2^-/-^* hosts was magnetically depleted of CD3^+^, CD8^+^, CD25^+^, B220^+^, Gr-1^+^, CD11b^+^, Ter119^+^ cells using Rat anti-mouse antibodies (BioXcell, NH, USA) and Dynabeads™ sheep anti-rat IgG beads (Invitrogen, MA, USA) according to manufacturer’s recommendations, then 10^7^ cells were injected into lethally irradiated *β2m^-/-^* hosts. The bone marrow chimera recipients were maintained for 4-6 weeks before collecting OT-I pre-selection thymocytes. For pre-positive selection OT-II thymocytes, OT-II *Rag2^-/-^ MHCII^-/-^* mice were bred. Thymic single cell suspensions from the OT-I *Rag2^-/-^* -> *β2m^-/-^* bone marrow chimeras or OT-II *Rag2^-/-^ MHCII^-/-^* mice were isolated, stained in 5mL RPMI 1640 + 5µM CellTrace™ Violet (Invitrogen) for 30 minutes in the dark at 37°C, washed once with warm complete RPMI, resuspended in fresh, warm complete RPMI, and kept in the dark at 37°C prior to slice addition. Prepared cells were overlaid on WT, *β2m^-/-^*, *MHCII^-/-^* or pCX-EGFP slices and incubated at 5% CO_2_ 37°C. Slices were gently washed at 3h with warm complete RPMI in a dish to prevent further entry of new cells. At the indicated time points, thymocytes were harvested by mechanical disruption of the slices for flow cytometry or intact slices were imaged by 2-photon microscopy.

### 2-photon imaging and analysis of thymic slices

For imaging of purified thymocyte subsets, sorted cells from each host were centrifuged for collection, then separately stained in RPMI 1640 with 5µM CellTracker™ Red CMTPX (Thermo Fisher Scientific, MA, USA) or Indo-1 AM (eBioscience, CA, USA) for 30 minutes in 37°C water bath, away from light. Dye used for each host was swapped between experiments to account for potential effect of dyes on cell viability and mobility. For imaging of pre-positive selection thymocytes, whole thymus single cells were stained in RPMI 1640 with 5µM CellTracker™ Red CMTPX in the same way. Stained cells were collected by centrifugation and resuspended in fresh warm cRPMI, incubated for 30 minutes in 37°C water bath, away from light. Prepared cells were collected by centrifugation, then washed 2 times with fresh warm complete RPMI to remove dye residues, then laid onto pCX-EGFP slices, and incubated in 5% CO_2_ incubator at 37°C for a minimum of 1 hour before 2-photon imaging.

For image and Video acquisition, thymic slices were secured to an imaging chamber (Harvard Apparatus, MA, USA) perfused with heated DRPMI (Corning) + 2g/L sodium bicarbonate, 5mM HEPES (Sigma-Aldrich, MA, USA) and 1.25mM CaCl_2_, pH = 7.4, aerated with 95% O_2_ + 5% CO_2_ at flow rate of 100mL/h. The imaging chamber is secured on a heated stage, and a temperature probe was inserted to monitor and maintain the chamber temperature at 37°C. Images were acquired using an Ultima IV microscope controlled by PrairieView software (V5.4, Bruker, MA, USA), with a 20X water immersion objective, NA = 1.0 (Olympus, Japan). Time-lapse Videos were acquired by a t-series of 60 rounds of 15s acquisition, through 40µm depth at 5µm intervals. Samples were illuminated with a Mai Tai Ti:sapphire laser (Spectra Physics) tuned to 865nm for simultaneous excitation of EGFP and CMTPX, and an additional InSight Ti:sapphire laser (Spectra Physics, CA, USA) tuned to 730nm for excitation of Indo-1 AM, if necessary. Emitted light was passed through 473/24, 525/50 and 605/70 band pass filters (Chroma, VT, USA) to separate PMTs for detecting Indo-1 AM (low calcium), EGFP and CMTPX signals, respectively.

Captured images were analyzed using Imaris v9.7.2 (Bitplane, Switzerland). For medullary enrichment analysis, medullary and cortical volumes were established by manually generating surface objects by morphological distinction. Fluorescently labelled cells were identified using Spot tools from random frames of each Video, and medullary or cortical cellular localization was determined by calculating the spot’s distance to the medullary surface object using the Spot distance to Surface function in ImarisXT.

Densities of cells within each region were calculated by dividing the number of cells in the respective region by the volume of the surface object, and medullary-to-cortical ratio was calculated by dividing the medullary density to the cortical density within the same Video, using Excel (Microsoft). For thymocyte migration parameters, trajectory of thymocytes were established with the automated Spot tracking feature within Imaris. Only trajectories with time span ≥ 3 minutes were included in the analysis. Average track speed and straightness were generated by Imaris for analysis.

### cDNA preparation and qPCR

FACS purified, frozen thymic APC subsets were thawed and lysed in TRIzol (Thermo Fisher), RNA was extracted, and cDNA waw synthesized using the qScript cDNA Synthesis Kit (QuantaBio, MA, USA), per manufacture’s recommendations. qRT-PCR was performed using SYBR Green PCR Master Mix (Applied Biosystems, MA, USA), on a ViiA 7 Real-Time PCR System (Applied Biosystems), using the following primers: *Ccl17* Forward: 5’-AGT GGA GTG TTC CAG GGA TG-3’, *Ccl17* Reverse: 5’-CCA ATC TGA TGG CCT TCT TC-3’, *Ccl19* Forward: 5’-GCT AAT GAT GCG GAA GAC TG-3’, *Ccl19* Reverse: 5’-ACT CAC ATC GAC TCT CTA GG-3’, *Ccl21* Forward: 5’-GCA GTG ATG GAG GGG GTC AG-3’, *Ccl21* Reverse: 5’-CGG GGT GAG AAC AGG ATT GC-3’, *Ccl22* Forward: 5’-AGG TCC CTA TGG TGC CAA TGT-3’, *Ccl22* Reverse: 5’- CGG CAG GAT TTT GAG GTC CA-3’, β-actin Forward: 5’- CAC TGT CGA GTC GCG TCC A-3’, β-actin Reverse: 5’- CAT CCA TGG CGA ACT GGT GG-3’. ddCT Relative expression levels of target genes in each subset were quantified by first normalizing to actin expression within each subset, then normalizing between subsets for expression of target genes relative to a subset previously shown to express high levels of that gene. For CCL17 and CCL22, expression levels were normalized to cDC2; For CCL19 and CCL21, expression levels were normalized to mTEC^lo^.

### Congenic TLR stimulation and co-culture experiment

To activate APCs, congenic CD45.1 mouse spleen was isolated and red blood cells were lysed with RBC lysis buffer (BioLegend, CA, USA). Leukocytes were then resuspended in complete RPMI with 1µg/mL high molecular weight Poly(I:C), 1µg/mL low molecular weight Poly(I:C), 100ng/mL LPS, 500nM ODN1826 (Invivogen, CA, USA), or without any TLR ligands, and incubated in 5% CO_2_ incubator at 37°C overnight. The next day, CD45.2 WT and *Ccr4^-/-^* CD4^+^ T cells were isolated from spleen by magnetically depleting CD8^+^, CD25^+^, B220^+^, CD11b^+^, Gr-1^+^ and Ter119^+^ cells with corresponding rat anti-mouse antibodies (BioXCell) and Dynabeads® sheep anti-rat IgG (Invitrogen), and resuspended in complete RPMI. Stimulated CD45.1 splenocytes were washed three times with fresh complete RPMI. 2.5×10^4^ WT or *Ccr4^-/-^* CD4^+^ purified T cells were plated with or without 1.25×10^5^ unstimulated or stimulated splenocytes per well in complete RPMI in a 96-well, U-bottom plate. At Day 3 and Day 5, each well was harvested and subjected to flow cytometry analysis. 15µm polyester beads of known quantities were added to quantify cells in each sample.

### Immunofluorescent analyses of thymic and LNs cryosections and detection of autoantibodies

To detect anti-nuclear autoantibodies in mouse serum, 7-µm cryosections were prepared from *Rag2^-/-^* kidneys (C57BL/6J background). Cryosections were fixed in acetone (#x02212;20°C for 20 minutes), rinsed 3X (2x PBS rinses and 1x PBS with 0.1% Tween 20), blocked with 9% donkey serum (Jackson ImmunoResearch) in PBS for 20 minutes and incubated with undiluted mouse serum from mice of the indicated genotypes for 2 hours at room temperature. After washing, auto-antibodies were detected by staining with anti-mouse IgG AF594 in PBS for 1 hour at room temperature. After washing, slides were incubated with DAPI, washed, and coverslips were mounted in ProLong^TM^ Gold Antifade reagent (ThermoFisher). For CCL22 immunofluorescent staining and inguinal lymph node immunostaining, 7-µm cryosections were prepared from the thymus of 1 month old C57BL/6J mice and from lymph nodes of 5–6-month-old mice, respectively. All slides were fixed and stained as above. Thymus sections were stained with anti-CCL22 AF439, anti-CD11c-biotin and anti-CD31-APC overnight at room temperature. Inguinal lymph nodes were stained with anti-CCL22, anti-CD11c -biotin, and anti-CD31 - APC overnight at room temperature. For both, after washing, anti-goat-Dylight 488 and streptavidin Alexa Fluor 594 secondary antibodies were incubated for 1 hour at room temperature. The DAPI nuclear stain was added and slides mounted as above. Immunofluorescent images were acquired on a DMi8 microscope (Leica), using a 10x/0.4 NA objective or a 20X/0.7NA objective. Stitched images were generated with LasX software (Leica). All images were uniformly processed and converted into Tiffs using Fiji software (ImageJ),

### H&E staining and quantification

Lacrimal glands, submandibular glands, colon, and liver were extracted from 6–8-month-old mice and fixed in formalin (Fisherbrand) for 48 hours before storing in 70% ethanol. Organs were sent to the Histology and Immunohistochemistry Laboratory at the University of Texas San Antonio for embedding, sectioning, and staining. Spleen and mLNs were extracted and processed by the Histology and Tissue Processing core at the University of Texas MD Anderson Cancer Center Science Park (Smithville, TX). Histological analysis of spleen, inguinal lymph nodes, liver and colon were done by a veterinary pathologist, while infiltrates in lacrimal and submandibular glands stains were analyzed in-house, as in previously described (Lieberman et al., 2015). Briefly, we scored foci composed of at least 50 mononuclear cells. In the cases where multiple foci coalesced, we assigned those foci a score value of 3 for statistical analysis. The number of inflammatory foci per 10 mm^2^ was calculated by counting the total number of foci by standard light microscopy using a 10 × objective and dividing that by the surface area of sections measured by Fiji software. Colon sections were scored for mucosal changes such as hyperplasia and architectural distortion, inflammation severity, and the percentage of the tissue affected as described previously (Hale et al., 2005), but with a single score that reflected the entire segment examined (total possible scores ranged from 0 – 15).

### Statistical analysis

All statistical analysis was conducted using Graphpad PRISM v9.8 (Graphpad Software, CA, USA), with the corresponding statistical tests and multiple comparison corrections listed in the figure legends.

## Acknowledgements

The authors thank the Animal Resource Center staff at the University of Texas at Austin for assistance with mouse maintenance, Dr. Jessica Lancaster for assistance with 2-photon microscopy training, Richard Salinas at the Center for Biomedical Research Support, and Dr. Ellen Richie for providing advice. Graphical illustrations were created with Biorender.com. This research was supported by a grant from the National Institutes of Health R01AI104870 to L.I.R.E.

## Author contributions

Y.L. and P.G.T. designed and conducted experiments, analyzed data, and wrote and edited the manuscript. J.S. and H.J.S. conducted experiments. L.P.H carried out histologic analyses and edited the manuscript. L.I.R.E. designed experiments, analyzed data, and edited the manuscript.

## Competing Interests statement

The authors have no conflicts of interest to declare.

## Supplemental Information

## Supplemental Figures and Legends

**Figure 1—figure supplement 1.**
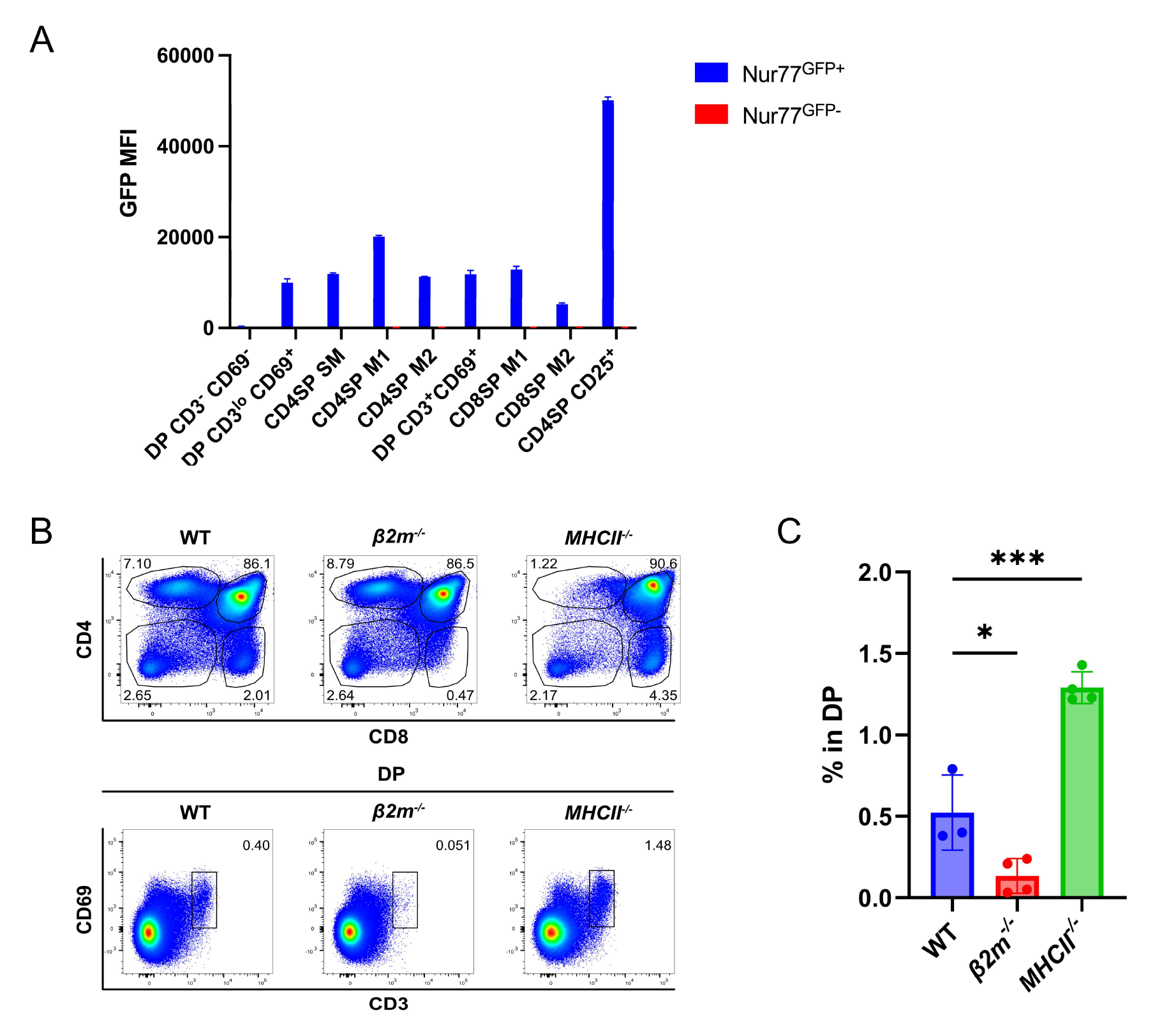
DP CD3^lo^CD69^+^ cells are consistent with post-positive selection DPs and DP CD3^+^CD69^+^ cells represent co-receptor reversing MHCI-restricted thymocytes. **A)** Quantification of the GFP MFI of the indicated thymocyte subsets from Nur77^GFP^ mice. Data were compiled from 3 independent experiments (Nur77^GFP+^ N=4 mice, Nur77^GFP-^ N=3 mice). **B)** Flow cytometry plots showing the absence of DP CD3^+^CD69^+^ thymocytes from *β2m^-/-^*, but not *MHCII^-/-^* thymus. **C)** Quantification of the frequency of DP CD3^+^CD69^+^ in all DP thymocytes from wild-type, *β2m^-/-^*, and *MHCII^-/-^* mice. One-way ANOVA with Dunnett’s multiple comparison correction was used for statistical analysis. (n=3 for wild-type, n=4 for *β2m^-/-^*, n=3 for *MHCII^-/-^*. Mean ± SEM; *: p<0.05, ***: p<0.001).

**Figure 1—figure supplement 2.**
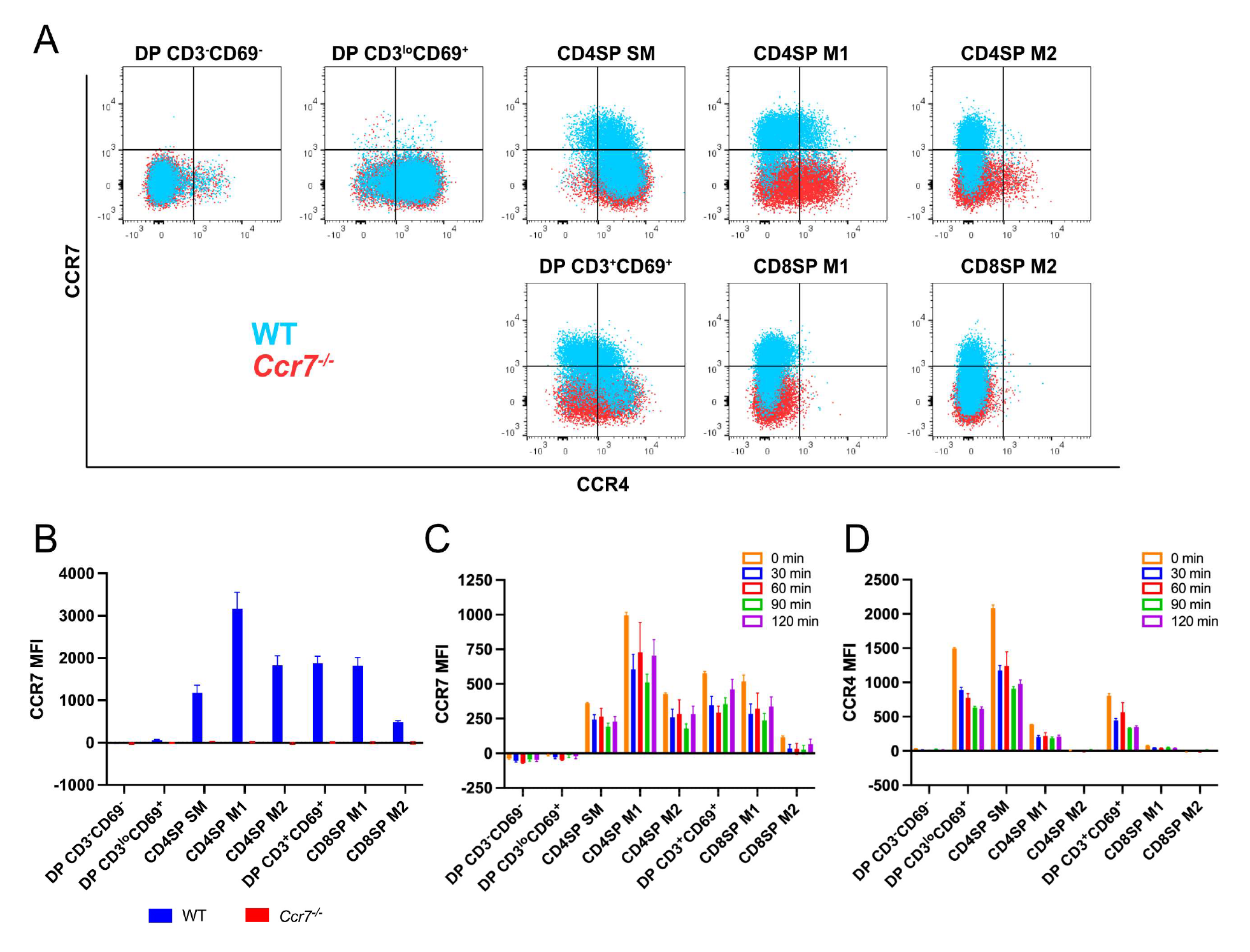
CCR7 expression on developing thymocytes begin at the CD4SP SM stage. **A)** Representative flow cytometry overlays showing CCR4 and CCR7 expression by post-positive selection thymocyte subsets from WT and *Ccr7^-/-^* mice, respectively. **B)** Quantification of the CCR7 MFI within each subset from data in A (N = 4 mice of each genotype). **C-D)** Quantification of CCR7 and CCR4 MFIs for each thymocyte subset after incubation at 37°C for the indicated times, prior to antibody staining at 4°C. Data were compiled from 2 independent experiments (n=4 biologic replicates).

**Figure 2—figure supplement 1.**
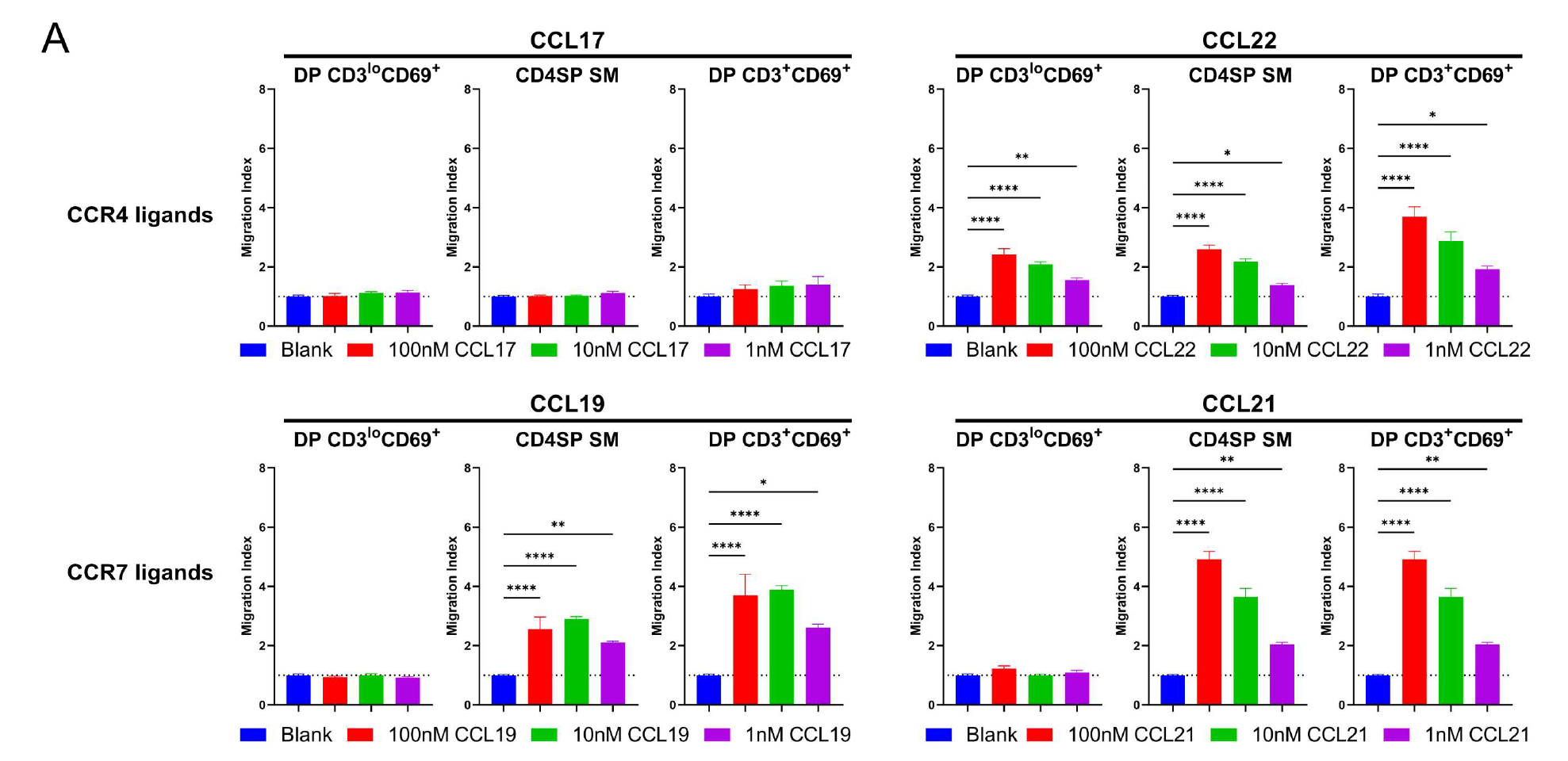
Chemokine signaling is not diminished in CD4SP SM thymocytes. **A)** Migration indices from data in Fig. 2 for DP CD3^lo^CD69^+^, CD4SP SM and DP CD3^lo^CD69^+^ were analyzed separately to compare the relative efficiency of CCR4 ligands and CCR7 ligands at inducing chemotaxis of each subset. Ordinary one-way ANOVA with Dunnett’s multiple comparison corrections were used for statistical analysis. (Mean ± SEM; *: p<0.05, **: p<0.01, ***: p<0.001, ****: p<0.0001)

**Figure 2—figure supplement 2.**
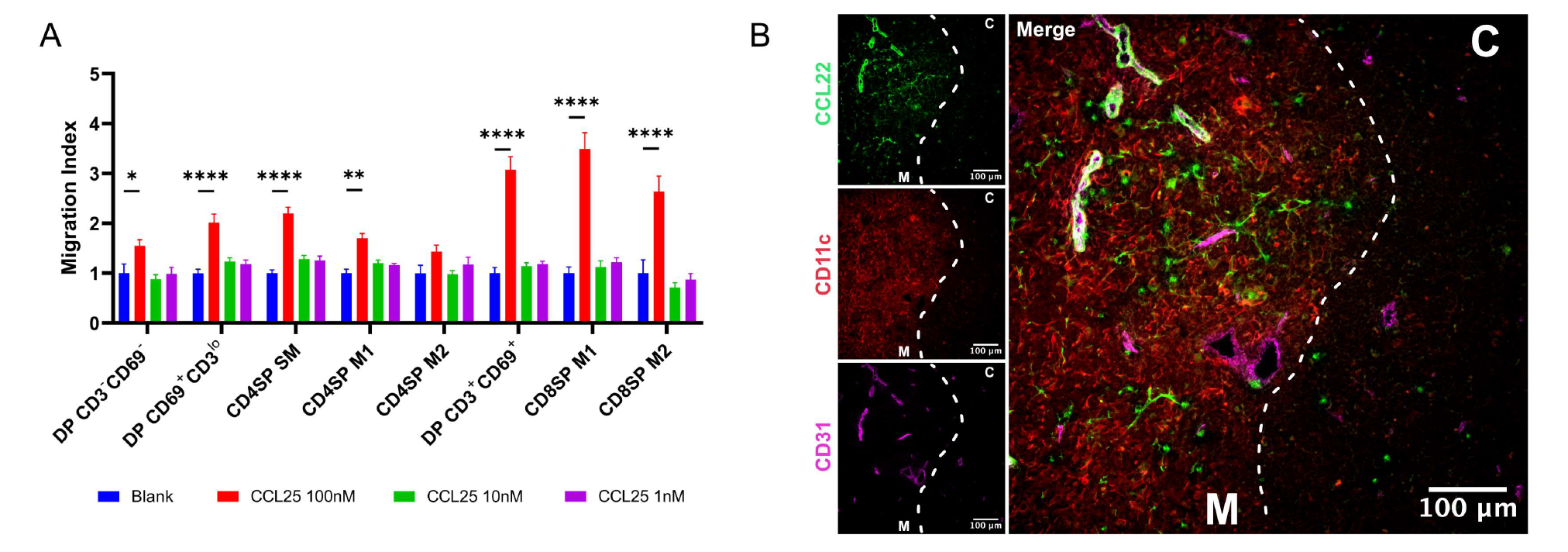
CD4SP SM thymocytes migrate towards CCL25 and CCL22 is localized in the medulla. **A)** Transwell assays were used to quantify chemotaxis of thymocyte subsets to the indicated concentrations of the CCR9 ligand, CCL25. Migration indexes were calculated as the frequencies of input cells of each subset that migrated towards the chemokine relative to the frequencies of input cells that migrated in the absence of chemokine (Blank). Data were compiled from 3 independent experiments with triplicate wells per assay. 2-way ANOVA with Dunnett’s multiple comparison correction were used for statistical analysis. (Mean ± SEM; *: p<0.05, **: p<0.01, ****: p<0.0001). **B)** Representative confocal images of CCL22 (green), CD11c (red), and CD31 (purple) on immunostained cryosections from WT thymus. The cortex (C) and medulla (M) are demarcated by white dotted lines.

**Figure 4— figure supplement 1.**
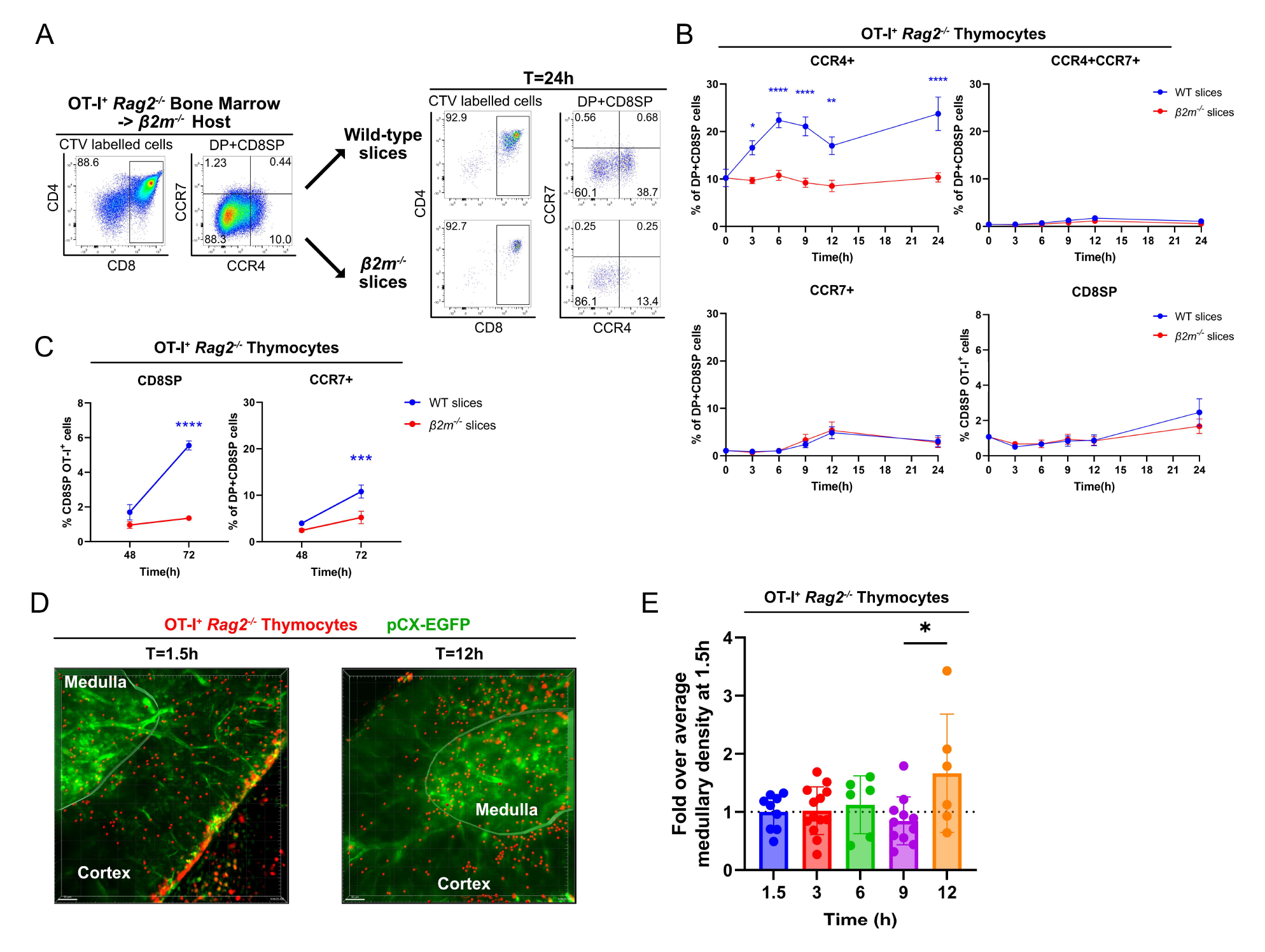
Rapid upregulation of CCR4 following positive selection of OT-I thymocytes correlates with medullary entry. **A)** Representative flow cytometry plots of CCR4 and CCR7 expression by pre-positive selection OT-I^+^ *Rag2^-/-^* thymocytes from *β2m^-/-^* hosts that were added to thymic slices (left) or analyzed 24h after incubation in WT or *β2m^-/-^*slices, as indicated (right). **B)** Percentages of CCR4^+^, CCR4^+^CCR7^+^, CCR7^+^ or CD8SP OT-I^+^ *Rag2^-/-^* thymocyte safter incubation on thymic slices of the indicated genotypes for the indicated time period up to 24h. **C)** Percentages of CCR7^+^ or CD8SP OT-I^+^ *Rag2*^-/-^ thymocytes at 48h and 72h time points after initiation of thymic slices assays. Data for B-C were compiled from 3 OT-I^+^ *Rag2^-/-^* BM -> *β2m^-/-^* experiments, with triplicate slices per experiment. Mixed-effect analysis with Šídák’s multiple comparison correction were used for statistical analysis. (Mean ± SEM; *: p<0.05, **: p<0.01, ****: p<0.0001) **D)** Representative maximum intensity projections of 2-photon imaging data showing a pCX-EGFP thymic slice (green) containing CMTPX-labeled OT-I^+^ *Rag2^-/-^* BM -> *β2m^-/-^* thymocytes (red) at 1.5h and 12h after thymocytes were added to the slices. Medullary and cortical volumes were demarcated as indicated by the masked regions. **E)** Quantification of medullary density of OT-I^+^ *Rag2^-/-^* cells at the indicated time points. Individual data points represent medullary density calculated from each Video, relative to the average medullary density of all Videos taken at 1.5 hour. Data were compiled from 3 time-course experiments. One-way ANOVA with Tucky’s multiple comparison correction were used for statistical analysis. (Mean ± SEM; *: p<0.05)

**Figure 5— figure supplement 1.**
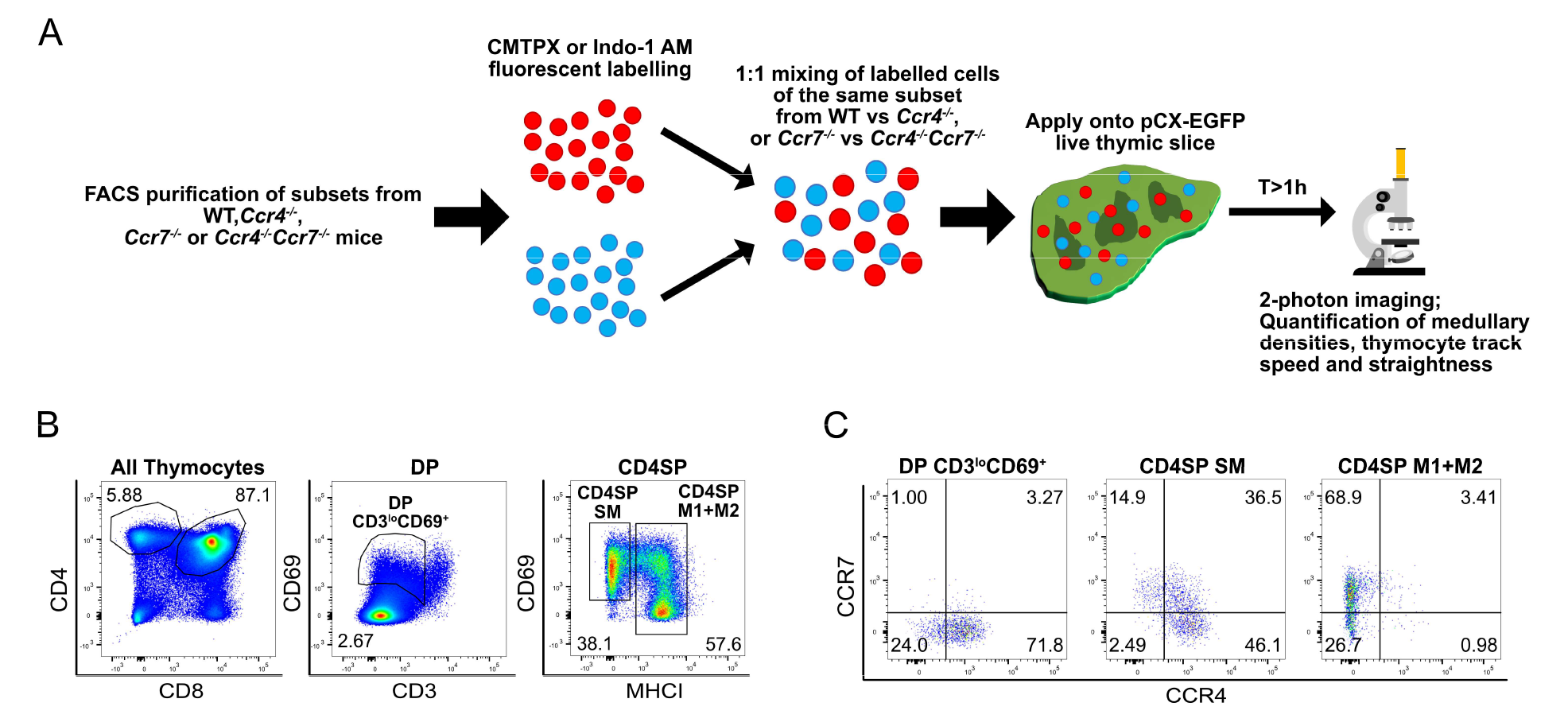
Experimental approach to test if CCR4 and CCR7 are required for medullary entry of distinct FACS-purified post-positive selection thymocyte subsets that differentially express CCR4 and CCR7. **A)** Experimental schematic of 2-photon imaging of sorted, fluorescently labeled post-positive selection thymocyte subsets. WT cells were imaged in the same slices with *Ccr4*^-/-^ or *Ccr7*^-/-^ cells, and *Ccr7*^-/-^ cells were imaged in the same slices with *Ccr4*^-/-^ *Ccr7*^-/-^ cells. Dyes were swapped between different experiments, and we did not find that use of CMTPX versus Indo-1 altered the results. **B-C)** Representative flow cytometry plots showing B) the gating scheme for sorting DP CD3^lo^CD69^+^, CD4SP SM and CD4SP M1+M2 subsets for 2-photon imaging, and C) CCR4 and CCR7 expression of the FACS purified thymocyte subsets from a WT mouse.

**Figure 6— figure supplement 1.**
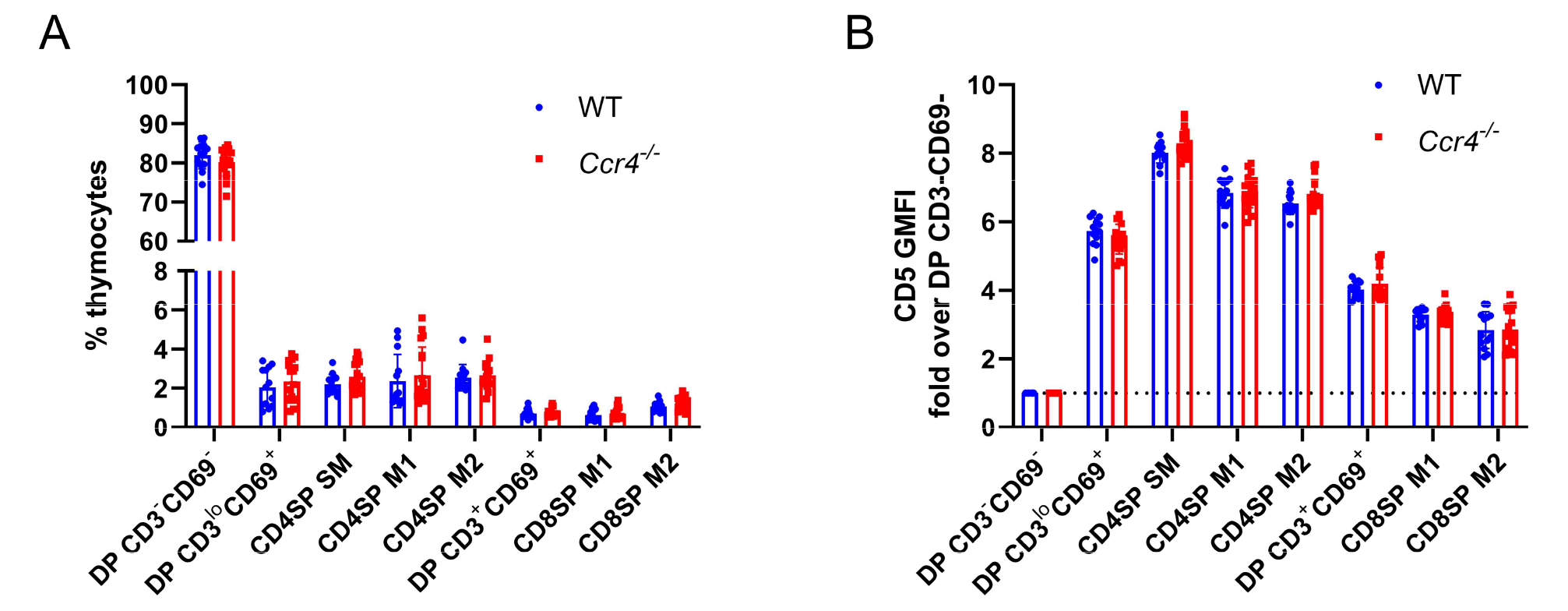
Thymocyte composition and CD5 levels are not altered in *Ccr4^-/-^* mice. **A)** Thymocyte subset frequencies in littermate WT and *Ccr4^-/-^*mice. **B)** CD5 expression levels on the indicated thymocyte subsets from littermate WT and *Ccr4^-/-^*mice, normalized to the DP CD3^-^CD69^-^ subset of each mouse. For A) and B), data were compiled from 6 independent experiments and significance determined using unpaired t-tests with Holm-Šídák’s multiple comparison correction (Mean ± SEM; n=13 for WT, n=18 for *Ccr4^-/-^*).

**Figure 7— figure supplement 1.**
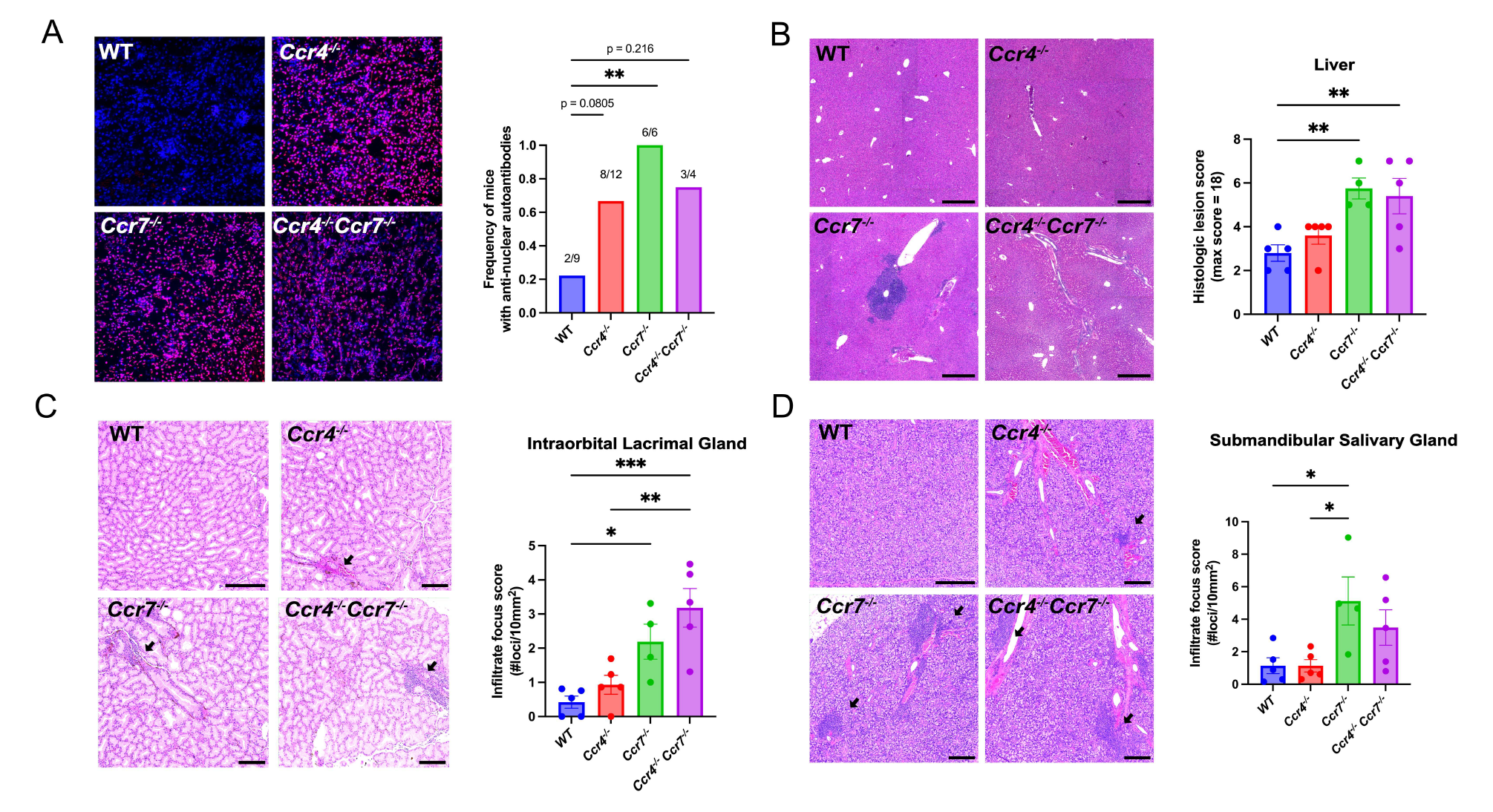
CCR4 and CCR7 support self-tolerance in different organs. **A)** Representative images of Rag2^-/-^ kidney sections immunostained with serum from 5–6.5-month-old *WT, Ccr4^-/-^, Ccr7^-/-^ or Ccr4^-/-^Ccr7^-/-^* mice (left). Anti-nuclear autoantibodies were detected as an overlay of anti–mouse IgG (red) and the nuclear DAPI stain (blue). Proportion of mice containing anti-nuclear serum autoantibodies was quantified (right). p < 0.05 **: p<0.01 (Fisher’s exact test)**. B)** Representative images of H&E stains from the liver of 5–6-month-old *WT, Ccr4^-/-^, Ccr7^-/-^* and *Ccr4^-/-^Ccr7^-/-^* mice (left). Bars, 500 µm. Histological lesion score was quantified by a veterinary pathologist (right). One-way ANOVA was used to test significance between groups (Mean ± SEM; **: p>0.01). **C-D)** Representative images of H&E stains from intraorbital lacrimal glands (C) and submandibular salivary glands (D) of 5–6- month-old *WT, Ccr4^-/-^, Ccr7^-/-^ or Ccr4^-/-^Ccr7^-/-^* mice (left). Bars, 500 µm. Arrows indicate lymphocytic infiltrate foci. The number of infiltrate foci per 10mm^2^ was quantified (right). One-way ANOVA was used to determine the significant differences between groups (Mean ± SEM; *: p<0.05, **: p<0.01, ***: p<0.001).

**Figure 7— figure supplement 2.**
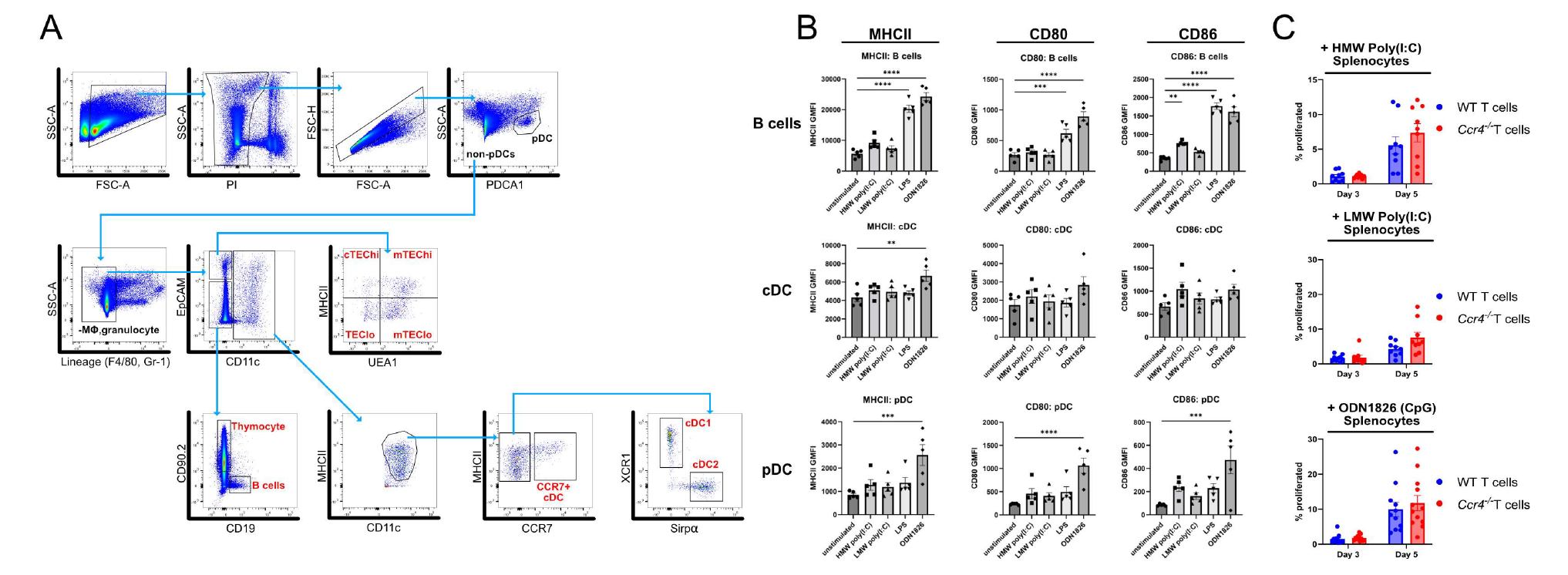
CCR4 promotes tolerance to activated antigen presenting cells. **A)** Gating scheme for FACS sorting of thymic APC subsets analyzed by qRT-PCR for expression of CCR4 and CCR7 ligands. **B)** Surface expression of MHCII, CD80 and CD86 on APC subsets from unstimulated or TLR ligand-stimulated splenocytes used in the proliferation assays in Figure 7F. Individual data points represent the GMFI of the indicated markers on each respective APC subset. Data were compiled from 5 individual experiments. One-way ANOVA with Dunnett’s multiple comparison correction was used for statistical analysis. (Mean ± SEM; **: p<0.01, ***: p<0.001 ****: p<0.0001) **C)** The percentage of WT and *Ccr4^-/-^* T cells that proliferated at days 3 and 5 when cultured with High Molecular Weight (HMW) poly(I:C), Low Molecular Weight (LMW) poly(I:C), or ODN1826 (CpG)-stimulated splenocytes. Data were compiled from 3-4 independent experiments with triplicate wells, and unpaired t-tests with Holm-Šídák’s multiple comparison correction was used for statistical analysis. (Mean ± SEM)

## Supplemental Video Legends

**Video S1.** Two-photon live imaging of OT-II+ *Rag2^-/-^ MHCII^-/-^* input thymocytes (red) migrating in a pCX-EGFP thymic slice (green) 1.5 hours after addition to positively selecting thymic slices, as illustrated in Figure 3D. Images were acquired for 15 minutes at 15 second time intervals through a depth of 40mm, and maximum intensity projections of input cells shown side by side.

**Video S2.** Two-photon live imaging of OT-II+ *Rag2^-/-^ MHCII^-/-^* input thymocytes (red) migrating in a pCX-EGFP thymic slice (green) 12 hours after addition to positively selecting thymic slices, as illustrated in Figure 3D. Images were acquired for 15 minutes at 15 second time intervals through a depth of 40mm, and maximum intensity projections of input cells shown side by side.

**Video S3.** Two-photon live imaging of OT-I+ *Rag2^-/-^*input thymocytes from a *β2m^-/-^*bone marrow chimera host (red) migrating in a pCX-EGFP thymic slice (green) 1.5 hours after addition to positively selecting thymic slices, as illustrated in Supplemental Figure 3D, respectively. Images were acquired for 15 minutes at 15 second time intervals through a depth of 40mm, and maximum intensity projections of input cells shown side by side.

**Video S4.** Two-photon live imaging of OT-I+ *Rag2^-/-^*input thymocytes from a *β2m^-/-^*bone marrow chimera host (red) migrating in a pCX-EGFP thymic slice (green) 12 hours after addition to positively selecting thymic slices, as illustrated in Supplemental Figure 3D, respectively. Images were acquired for 15 minutes at 15 second time intervals through a depth of 40mm, and maximum intensity projections of input cells shown side by side.

**Video S5.** Two-photon live imaging of WT (red) or *Ccr4^-/-^* (blue) FACS sorted DP CD3^lo^CD69^+^ thymocytes migrating in pCX-EGFP thymic slices (green), as illustrated in Figure 4A. Images were acquired for 15 minutes at 15 second time intervals through a depth of 40mm, and maximum intensity projections of input cells with and without cell tracks, color-coded for elapsed imaging time, are shown side by side.

**Video S6.** Two-photon live imaging of WT (red) or *Ccr4^-/-^*(blue) FACS sorted CD4SP SM thymocytes migrating in pCX-EGFP thymic slices (green), as illustrated in Figure 4A. Images were acquired for 15 minutes at 15 second time intervals through a depth of 40mm, and maximum intensity projections of input cells with and without cell tracks, color-coded for elapsed imaging time, are shown side by side.

**Video S7.** Two-photon live imaging of WT (red) or *Ccr4^-/-^*(blue) FACS sorted CD4SP M1+M2 thymocyte subsets, migrating in pCX-EGFP thymic slices (green), as illustrated in Figure 4A. Images were acquired for 15 minutes at 15 second time intervals through a depth of 40mm, and maximum intensity projections of input cells with and without cell tracks, color-coded for elapsed imaging time, are shown side by side.

**Video S8.** Two-photon live imaging of *Ccr4^-/-^Ccr7^-/-^* (red) or *Ccr7^-/-^* (blue) FACS sorted CD4SP M1+M2 subsets, migrating in pCX-EGFP thymic slices (green). Images were acquired for 15 minutes at 15 second time intervals through a depth of 40mm, and maximum intensity projections of input cells with and without cell tracks, color-coded for elapsed imaging time, are shown side by side.

## Notes

### Competing Interest Statement

The authors have declared no competing interest.

### Summary of Updates

New 2-photon imaging data and flow cytometry analyses confirm and extend the requirement for CCR4 versus CCR7 in medullary accumulation and central tolerance of early versus late stage negative selection. Histologic evaluation reveals CCR4 and CCR7 are required to prevent autoimmunity/autoinflammation in different organs

## References

Abramson, J., & Anderson, G. (2017). Thymic Epithelial Cells. Annu Rev Immunol, 35(1), 85–118. https://doi.org/10.1146/annurev-immunol-051116-052320

Alam, A., Braun, M. Y., Hartgers, F., Lesage, S., Cohen, L., Hugo, P., Denis, F., & Sékaly, R.-P. (1997). Specific Activation of the Cysteine Protease CPP32 during the Negative Selection of T Cells in the Thymus. Journal of Experimental Medicine, 186(9), 1503-1512. https://doi.org/10.1084/jem.186.9.1503

Anderson, M. S., & Su, M. A. (2016). AIRE expands: new roles in immune tolerance and beyond. Nat Rev Immunol, 16(4), 247–258. https://doi.org/10.1038/nri.2016.9

Anderson, M. S., Venanzi, E. S., Klein, L., Chen, Z., Berzins, S. P., Turley, S. J., von Boehmer, H., Bronson, R., Dierich, A., Benoist, C., & Mathis, D. (2002). Projection of an immunological self shadow within the thymus by the aire protein. Science, 298(5597), 1395–1401. https://doi.org/10.1126/science.1075958

Ardouin, L., Luche, H., Chelbi, R., Carpentier, S., Shawket, A., Montanana Sanchis, F., Santa Maria, C., Grenot, P., Alexandre, Y., Gregoire, C., Fries, A., Vu Manh, T. P., Tamoutounour, S., Crozat, K., Tomasello, E., Jorquera, A., Fossum, E., Bogen, B., Azukizawa, H., . . . Malissen, B. (2016). Broad and Largely Concordant Molecular Changes Characterize Tolerogenic and Immunogenic Dendritic Cell Maturation in Thymus and Periphery. Immunity, 45(2), 305–318. https://doi.org/10.1016/j.immuni.2016.07.019

Atibalentja, D. F., Byersdorfer, C. A., & Unanue, E. R. (2009). Thymus-blood protein interactions are highly effective in negative selection and regulatory T cell induction. J Immunol, 183(12), 7909–7918. https://doi.org/10.4049/jimmunol.0902632

Au-Yeung, B. B., Melichar, H. J., Ross, J. O., Cheng, D. A., Zikherman, J., Shokat, K. M., Robey, E. A., & Weiss, A. (2014). Quantitative and temporal requirements revealed for Zap70 catalytic activity during T cell development. Nat Immunol, 15(7), 687–694. https://doi.org/10.1038/ni.2918

Azzam, H. S., Grinberg, A., Lui, K., Shen, H., Shores, E. W., & Love, P. E. (1998). CD5 expression is developmentally regulated by T cell receptor (TCR) signals and TCR avidity. J Exp Med, 188(12), 2301–2311. https://doi.org/10.1084/jem.188.12.2301

Bendelac, A., Killeen, N., Littman, D. R., & Schwartz, R. H. (1994). A subset of CD4+ thymocytes selected by MHC class I molecules. Science, 263(5154), 1774–1778. https://doi.org/10.1126/science.7907820

Bleul, C. C., & Boehm, T. (2000). Chemokines define distinct microenvironments in the developing thymus. Eur J Immunol, 30(12), 3371–3379. https://doi.org/10.1002/1521-4141(2000012)30:12<3371::AID-IMMU3371>3.0.CO;2-L

Bonasio, R., Scimone, M. L., Schaerli, P., Grabie, N., Lichtman, A. H., & von Andrian, U. H. (2006). Clonal deletion of thymocytes by circulating dendritic cells homing to the thymus. Nat Immunol, 7(10), 1092–1100. https://doi.org/10.1038/ni1385

Bornstein, C., Nevo, S., Giladi, A., Kadouri, N., Pouzolles, M., Gerbe, F., David, E., Machado, A., Chuprin, A., Toth, B., Goldberg, O., Itzkovitz, S., Taylor, N., Jay, P., Zimmermann, V. S., Abramson, J., & Amit, I. (2018). Single-cell mapping of the thymic stroma identifies IL-25-producing tuft epithelial cells. Nature, 559(7715), 622–626. https://doi.org/10.1038/s41586-018-0346-1

Boursalian, T. E., Golob, J., Soper, D. M., Cooper, C. J., & Fink, P. J. (2004). Continued maturation of thymic emigrants in the periphery. Nat Immunol, 5(4), 418–425. https://doi.org/10.1038/ni1049

Breed, E. R., Watanabe, M., & Hogquist, K. A. (2019). Measuring Thymic Clonal Deletion at the Population Level. J Immunol, 202(11), 3226–3233. https://doi.org/10.4049/jimmunol.1900191

Brennecke, P., Reyes, A., Pinto, S., Rattay, K., Nguyen, M., Kuchler, R., Huber, W., Kyewski, B., & Steinmetz, L. M. (2015). Single-cell transcriptome analysis reveals coordinated ectopic gene-expression patterns in medullary thymic epithelial cells. Nat Immunol, 16(9), 933–941. https://doi.org/10.1038/ni.3246

Britschgi, M. R., Link, A., Lissandrin, T. K. A., & Luther, S. A. (2008). Dynamic Modulation of CCR7 Expression and Function on Naive T Lymphocytes In Vivo1. The Journal of Immunology, 181(11), 7681–7688. https://doi.org/10.4049/jimmunol.181.11.7681

Brugnera, E., Bhandoola, A., Cibotti, R., Yu, Q., Guinter, T. I., Yamashita, Y., Sharrow, S. O., & Singer, A. (2000). Coreceptor reversal in the thymus: signaled CD4+8+ thymocytes initially terminate CD8 transcription even when differentiating into CD8+ T cells. Immunity, 13(1), 59–71. https://doi.org/10.1016/s1074-7613(00)00008-x

Campbell, J. J., Pan, J., & Butcher, E. C. (1999). Cutting edge: developmental switches in chemokine responses during T cell maturation. J Immunol, 163(5), 2353–2357. https://doi.org/ji_v163n5p2353 [pii]

Cepeda, S., Cantu, C., Orozco, S., Xiao, Y., Brown, Z., Semwal, M. K., Venables, T., Anderson, M. S., & Griffith, A. V. (2018). Age-Associated Decline in Thymic B Cell Expression of Aire and Aire-Dependent Self-Antigens. Cell Rep, 22(5), 1276–1287. https://doi.org/10.1016/j.celrep.2018.01.015

Chvatchko, Y., Hoogewerf, A. J., Meyer, A., Alouani, S., Juillard, P., Buser, R., Conquet, F., Proudfoot, A. E. I., Wells, T. N. C., & Power, C. A. (2000). A Key Role for Cc Chemokine Receptor 4 in Lipopolysaccharide-Induced Endotoxic Shock. Journal of Experimental Medicine, 191(10), 1755–1764. https://doi.org/10.1084/jem.191.10.1755

Cowan, J. E., McCarthy, N. I., Parnell, S. M., White, A. J., Bacon, A., Serge, A., Irla, M., Lane, P. J., Jenkinson, E. J., Jenkinson, W. E., & Anderson, G. (2014). Differential requirement for CCR4 and CCR7 during the development of innate and adaptive alphabetaT cells in the adult thymus. J Immunol, 193(3), 1204–1212. https://doi.org/10.4049/jimmunol.1400993

Daley, S. R., Hu, D. Y., & Goodnow, C. C. (2013). Helios marks strongly autoreactive CD4+ T cells in two major waves of thymic deletion distinguished by induction of PD-1 or NF-kappaB. J Exp Med, 210(2), 269–285. https://doi.org/10.1084/jem.20121458

DeVoss, J., Hou, Y., Johannes, K., Lu, W., Liou, G. I., Rinn, J., Chang, H., Caspi, R. R., Fong, L., & Anderson, M. S. (2006). Spontaneous autoimmunity prevented by thymic expression of a single self-antigen. J Exp Med, 203(12), 2727–2735. https://doi.org/10.1084/jem.20061864

Dzhagalov, I. L., Chen, K. G., Herzmark, P., & Robey, E. A. (2013). Elimination of self-reactive T cells in the thymus: a timeline for negative selection. PLoS Biol, 11(5), e1001566. https://doi.org/10.1371/journal.pbio.1001566

Ehrlich, L. I., Oh, D. Y., Weissman, I. L., & Lewis, R. S. (2009). Differential contribution of chemotaxis and substrate restriction to segregation of immature and mature thymocytes. Immunity, 31(6), 986–998. https://doi.org/10.1016/j.immuni.2009.09.020

Control of Migration during Intrathymic T Cell Development, 1 Encyclopedia of Immunobiology 249-262 (Elsevier 2016).

Finnish-German, A. C. (1997). An autoimmune disease, APECED, caused by mutations in a novel gene featuring two PHD-type zinc-finger domains. Nat Genet, 17(4), 399–403. https://doi.org/10.1038/ng1297-399

Fu, G., Vallee, S., Rybakin, V., McGuire, M. V., Ampudia, J., Brockmeyer, C., Salek, M., Fallen, P. R., Hoerter, J. A., Munshi, A., Huang, Y. H., Hu, J., Fox, H. S., Sauer, K., Acuto, O., & Gascoigne, N. R. (2009). Themis controls thymocyte selection through regulation of T cell antigen receptor-mediated signaling. Nat Immunol, 10(8), 848–856. https://doi.org/10.1038/ni.1766

Hale, L. P., Greer, P. K., Trinh, C. T., & Gottfried, M. R. (2005). Treatment with oral bromelain decreases colonic inflammation in the IL-10-deficient murine model of inflammatory bowel disease. Clinical Immunology, 116(2), 135–142. https://doi.org/https://doi.org/10.1016/j.clim.2005.04.011

Hawiger, D., Masilamani, R. F., Bettelli, E., Kuchroo, V. K., & Nussenzweig, M. C. (2004). Immunological unresponsiveness characterized by increased expression of CD5 on peripheral T cells induced by dendritic cells in vivo. Immunity, 20(6), 695–705. https://doi.org/10.1016/j.immuni.2004.05.002

Hu, D. Y., Yap, J. Y., Wirasinha, R. C., Howard, D. R., Goodnow, C. C., & Daley, S. R. (2016). A timeline demarcating two waves of clonal deletion and Foxp3 upregulation during thymocyte development. Immunol Cell Biol, 94(4), 357–366. https://doi.org/10.1038/icb.2015.95

Hu, Z., Lancaster, J. N., & Ehrlich, L. I. (2015). The Contribution of Chemokines and Migration to the Induction of Central Tolerance in the Thymus. Front Immunol, 6, 398. https://doi.org/10.3389/fimmu.2015.00398

Hu, Z., Lancaster, J. N., Sasiponganan, C., & Ehrlich, L. I. (2015). CCR4 promotes medullary entry and thymocyte-dendritic cell interactions required for central tolerance. J Exp Med, 212(11), 1947–1965. https://doi.org/10.1084/jem.20150178

Hu, Z., Li, Y., Van Nieuwenhuijze, A., Selden, H. J., Jarrett, A. M., Sorace, A. G., Yankeelov, T. E., Liston, A., & Ehrlich, L. I. R. (2017). CCR7 Modulates the Generation of Thymic Regulatory T Cells by Altering the Composition of the Thymic Dendritic Cell Compartment. Cell Rep, 21(1), 168–180. https://doi.org/10.1016/j.celrep.2017.09.016

Kadakia, T., Tai, X., Kruhlak, M., Wisniewski, J., Hwang, I. Y., Roy, S., Guinter, T. I., Alag, A., Kehrl, J. H., Zhuang, Y., & Singer, A. (2019). E-protein-regulated expression of CXCR4 adheres preselection thymocytes to the thymic cortex. J Exp Med, 216(8), 1749–1761. https://doi.org/10.1084/jem.20182285

Ki, S., Park, D., Selden, H. J., Seita, J., Chung, H., Kim, J., Iyer, V. R., & Ehrlich, L. I. R. (2014). Global transcriptional profiling reveals distinct functions of thymic stromal subsets and age-related changes during thymic involution. Cell Rep, 9(1), 402–415. https://doi.org/10.1016/j.celrep.2014.08.070

Klein, L., Kyewski, B., Allen, P. M., & Hogquist, K. A. (2014). Positive and negative selection of the T cell repertoire: what thymocytes see (and don’t see). Nat Rev Immunol, 14(6), 377–391. https://doi.org/10.1038/nri3667

Klein, L., Robey, E. A., & Hsieh, C. S. (2019). Central CD4(+) T cell tolerance: deletion versus regulatory T cell differentiation. Nat Rev Immunol, 19(1), 7–18. https://doi.org/10.1038/s41577-018-0083-6

Koble, C., & Kyewski, B. (2009). The thymic medulla: a unique microenvironment for intercellular self-antigen transfer. J Exp Med, 206(7), 1505–1513. https://doi.org/10.1084/jem.20082449

Kozai, M., Kubo, Y., Katakai, T., Kondo, H., Kiyonari, H., Schaeuble, K., Luther, S. A., Ishimaru, N., Ohigashi, I., & Takahama, Y. (2017). Essential role of CCL21 in establishment of central self-tolerance in T cells. J Exp Med, 214(7), 1925–1935. https://doi.org/10.1084/jem.20161864

Kurobe, H., Liu, C., Ueno, T., Saito, F., Ohigashi, I., Seach, N., Arakaki, R., Hayashi, Y., Kitagawa, T., Lipp, M., Boyd, R. L., & Takahama, Y. (2006). CCR7-dependent cortex-to-medulla migration of positively selected thymocytes is essential for establishing central tolerance. Immunity, 24(2), 165–177. https://doi.org/10.1016/j.immuni.2005.12.011

Lancaster, J. N., & Ehrlich, L. I. R. (2017). Analysis of Thymocyte Migration, Cellular Interactions, and Activation by Multiphoton Fluorescence Microscopy of Live Thymic Slices. In G. E. Rainger & H. M. McGettrick (Eds.), T-Cell Trafficking: Methods and Protocols (pp. 9–25). Springer New York. https://doi.org/10.1007/978-1-4939-6931-9_2

Lancaster, J. N., Li, Y., & Ehrlich, L. I. R. (2018). Chemokine-Mediated Choreography of Thymocyte Development and Selection. Trends Immunol, 39(2), 86–98. https://doi.org/10.1016/j.it.2017.10.007

Lancaster, J. N., Thyagarajan, H. M., Srinivasan, J., Li, Y., Hu, Z., & Ehrlich, L. I. R. (2019). Live-cell imaging reveals the relative contributions of antigen-presenting cell subsets to thymic central tolerance. Nat Commun, 10(1), 2220. https://doi.org/10.1038/s41467-019-09727-4

Liu, B., Lin, Y., Yan, J., Yao, J., Liu, D., Ma, W., Wang, J., Liu, W., Wang, C., Zhang, L., & Qi, H. (2021). Affinity-coupled CCL22 promotes positive selection in germinal centres. Nature, 592(7852), 133–137. https://doi.org/10.1038/s41586-021-03239-2

Lutes, L. K., Steier, Z., McIntyre, L. L., Pandey, S., Kaminski, J., Hoover, A. R., Ariotti, S., Streets, A., Yosef, N., & Robey, E. A. (2021). T cell self-reactivity during thymic development dictates the timing of positive selection. eLife, 10, 1–28. https://doi.org/10.7554/eLife.65435

Marchetti, M. C., Di Marco, B., Cifone, G., Migliorati, G., & Riccardi, C. (2003). Dexamethasone-induced apoptosis of thymocytes: role of glucocorticoid receptor-associated Src kinase and caspase-8 activation. Blood, 101(2), 585–593. https://doi.org/10.1182/blood-2002-06-1779

Mathis, D., & Benoist, C. (2007). A decade of AIRE. Nat Rev Immunol, 7(8), 645–650. https://doi.org/10.1038/nri2136

McCaughtry, T. M., Baldwin, T. A., Wilken, M. S., & Hogquist, K. A. (2008). Clonal deletion of thymocytes can occur in the cortex with no involvement of the medulla. J Exp Med, 205(11), 2575–2584. https://doi.org/10.1084/jem.20080866

McCaughtry, T. M., Wilken, M. S., & Hogquist, K. A. (2007). Thymic emigration revisited. J Exp Med, 204(11), 2513–2520. https://doi.org/10.1084/jem.20070601

Melichar, H. J., Ross, J. O., Herzmark, P., Hogquist, K. A., & Robey, E. A. (2013). Distinct temporal patterns of T cell receptor signaling during positive versus negative selection in situ. Sci Signal, 6(297), ra92. https://doi.org/10.1126/scisignal.2004400

Meredith, M., Zemmour, D., Mathis, D., & Benoist, C. (2015). Aire controls gene expression in the thymic epithelium with ordered stochasticity. Nat Immunol, 16(9), 942–949. https://doi.org/10.1038/ni.3247

Misslitz, A., Pabst, O., Hintzen, G., Ohl, L., Kremmer, E., Petrie, H. T., & Forster, R. (2004). Thymic T cell development and progenitor localization depend on CCR7. J Exp Med, 200(4), 481–491. https://doi.org/10.1084/jem.20040383

Nagamine, K., Peterson, P., Scott, H. S., Kudoh, J., Minoshima, S., Heino, M., Krohn, K. J., Lalioti, M. D., Mullis, P. E., Antonarakis, S. E., Kawasaki, K., Asakawa, S., Ito, F., & Shimizu, N. (1997). Positional cloning of the APECED gene. Nat Genet, 17(4), 393–398. https://doi.org/10.1038/ng1297-393

Nitta, T., Nitta, S., Lei, Y., Lipp, M., & Takahama, Y. (2009). CCR7-mediated migration of developing thymocytes to the medulla is essential for negative selection to tissue-restricted antigens. Proc Natl Acad Sci U S A, 106(40), 17129–17133. https://doi.org/10.1073/pnas.0906956106

Oh, J., Wu, N., Barczak, A. J., Barbeau, R., Erle, D. J., & Shin, J. S. (2018). CD40 Mediates Maturation of Thymic Dendritic Cells Driven by Self-Reactive CD4(+) Thymocytes and Supports Development of Natural Regulatory T Cells. J Immunol, 200(4), 1399–1412. https://doi.org/10.4049/jimmunol.1700768

Perera, J., Zheng, Z., Li, S., Gudjonson, H., Kalinina, O., Benichou, J. I. C., Block, K. E., Louzoun, Y., Yin, D., Chong, A. S., Dinner, A. R., Weigert, M., & Huang, H. (2016). Self-Antigen-Driven Thymic B Cell Class Switching Promotes T Cell Central Tolerance. Cell Rep, 17(2), 387–398. https://doi.org/10.1016/j.celrep.2016.09.011

Persaud, S. P., Parker, C. R., Lo, W. L., Weber, K. S., & Allen, P. M. (2014). Intrinsic CD4+ T cell sensitivity and response to a pathogen are set and sustained by avidity for thymic and peripheral complexes of self peptide and MHC. Nat Immunol, 15(3), 266–274. https://doi.org/10.1038/ni.2822

Petrie, H. T., & Zuniga-Pflucker, J. C. (2007). Zoned out: functional mapping of stromal signaling microenvironments in the thymus. Annu Rev Immunol, 25, 649–679. https://doi.org/10.1146/annurev.immunol.23.021704.115715

Ross, J. O., Melichar, H. J., Au-Yeung, B. B., Herzmark, P., Weiss, A., & Robey, E. A. (2014). Distinct phases in the positive selection of CD8+ T cells distinguished by intrathymic migration and T-cell receptor signaling patterns. Proc Natl Acad Sci U S A, 111(25), E2550–2558. https://doi.org/10.1073/pnas.1408482111

Ruscher, R., Kummer, R. L., Lee, Y. J., Jameson, S. C., & Hogquist, K. A. (2017). CD8alphaalpha intraepithelial lymphocytes arise from two main thymic precursors. Nat Immunol, 18(7), 771–779. https://doi.org/10.1038/ni.3751

Sansom, S. N., Shikama-Dorn, N., Zhanybekova, S., Nusspaumer, G., Macaulay, I. C., Deadman, M. E., Heger, A., Ponting, C. P., & Hollander, G. A. (2014). Population and single-cell genomics reveal the Aire dependency, relief from Polycomb silencing, and distribution of self-antigen expression in thymic epithelia. Genome Res, 24(12), 1918–1931. https://doi.org/10.1101/gr.171645.113

Sinclair, C., Bains, I., Yates, A. J., & Seddon, B. (2013). Asymmetric thymocyte death underlies the CD4:CD8 T-cell ratio in the adaptive immune system. Proc Natl Acad Sci U S A, 110(31), E2905–2914. https://doi.org/10.1073/pnas.1304859110

Stritesky, G. L., Xing, Y., Erickson, J. R., Kalekar, L. A., Wang, X., Mueller, D. L., Jameson, S. C., & Hogquist, K. A. (2013). Murine thymic selection quantified using a unique method to capture deleted T cells. Proc Natl Acad Sci U S A, 110(12), 4679–4684. https://doi.org/10.1073/pnas.1217532110

Ueno, T., Hara, K., Willis, M. S., Malin, M. A., Hopken, U. E., Gray, D. H., Matsushima, K., Lipp, M., Springer, T. A., Boyd, R. L., Yoshie, O., & Takahama, Y. (2002). Role for CCR7 ligands in the emigration of newly generated T lymphocytes from the neonatal thymus. Immunity, 16(2), 205–218. https://doi.org/10.1016/s1074-7613(02)00267-4

Ueno, T., Saito, F., Gray, D. H., Kuse, S., Hieshima, K., Nakano, H., Kakiuchi, T., Lipp, M., Boyd, R. L., & Takahama, Y. (2004). CCR7 signals are essential for cortex-medulla migration of developing thymocytes. J Exp Med, 200(4), 493–505. https://doi.org/10.1084/jem.20040643

Vollmann, E. H., Rattay, K., Barreiro, O., Thiriot, A., Fuhlbrigge, R. A., Vrbanac, V., Kim, K. W., Jung, S., Tager, A. M., & von Andrian, U. H. (2021). Specialized transendothelial dendritic cells mediate thymic T-cell selection against blood-borne macromolecules. Nat Commun, 12(1), 6230. https://doi.org/10.1038/s41467-021-26446-x

Wright, D. E., Cheshier, S. H., Wagers, A. J., Randall, T. D., Christensen, J. L., & Weissman, I. L. (2001). Cyclophosphamide/granulocyte colony-stimulating factor causes selective mobilization of bone marrow hematopoietic stem cells into the blood after M phase of the cell cycle. Blood, 97(8), 2278–2285. https://doi.org/10.1182/blood.v97.8.2278

Xing, Y., Wang, X., Jameson, S. C., & Hogquist, K. A. (2016). Late stages of T cell maturation in the thymus involve NF-kappaB and tonic type I interferon signaling. Nat Immunol, 17(5), 565–573. https://doi.org/10.1038/ni.3419

Yamano, T., Nedjic, J., Hinterberger, M., Steinert, M., Koser, S., Pinto, S., Gerdes, N., Lutgens, E., Ishimaru, N., Busslinger, M., Brors, B., Kyewski, B., & Klein, L. (2015). Thymic B Cells Are Licensed to Present Self Antigens for Central T Cell Tolerance Induction. Immunity, 42(6), 1048–1061. https://doi.org/10.1016/j.immuni.2015.05.013

Zegarra-Ruiz, D. F., Kim, D. V., Norwood, K., Kim, M., Wu, W. H., Saldana-Morales, F. B., Hill, A. A., Majumdar, S., Orozco, S., Bell, R., Round, J. L., Longman, R. S., Egawa, T., Bettini, M. L., & Diehl, G. E. (2021). Thymic development of gut-microbiota-specific T cells. Nature, 594(7863), 413–417. https://doi.org/10.1038/s41586-021-03531-1

